# CD117 epitope-shielded hematopoietic stem cell transplantation with toxin-free conditioning and in vivo selection ameliorates a β-thalassemia model

**DOI:** 10.64898/2026.07.07.736903

**Authors:** Romina Marone, Rosalba Lepore, Kiriaki Paschoudi, Jessica Zuin, Alessandro Sinopoli, Anna Camus, Thomas Burgold, Ewelina Bartoszek, Diego Calabrese, Mylène Toranelli, Julia Wittwer, Manuel Rhiel, Geoffroy Andrieux, Chang Li, Alvin Hsu, Amélie Wiederkehr, Lisa C. Wellinger, Eva-Maria Grossjohann, Emiel Ten Buren, Julie Brault, Laura Garcia Prat, Frank Lehmann, Valentin Do Sacramento, Christopher Divsalar, Saniye Yumlu, David R. Liu, André Lieber, Toni Cathomen, Tatjana I. Cornu, Evangelia Yannaki, Stefanie Urlinger, Lukas T. Jeker

**Author notes:** Contributed equally (Rosalba Lepore, Kiriaki Paschoudi, Jessica Zuin, Alessandro Sinopoli). corresponding **Author Correspondence:** Lukas T. Jeker, MD, PhD Department of Biomedicine, Basel University Hospital and University of Basel, Hebelstrasse 20, CH-4031 Basel, Switzerland Or Romina Marone, PhD Department of Biomedicine, Basel University Hospital and University of Basel, Hebelstrasse 20, CH-4031 Basel, Switzerland.

## Abstract

Clinical evidence demonstrates that ex vivo gene therapy and genome engineering of hematopoietic stem and progenitor cells (HSPCs) could represent one-time cures. However, while genome editing itself has become increasingly efficient and precise, the toxic conditioning required for hematopoietic stem cell transplantation remains a major barrier to broad clinical implementation of these otherwise curative therapies. In particular, the use of busulfan for myeloablative conditioning constitutes a major safety concern. While preclinical studies established CD117 as a promising target for antigen-specific therapy, clinical translation faced setbacks balancing efficacy and safety. To overcome current limitations, we generated a new CD117-blocking monoclonal antibody (CIM058) and demonstrate its potency to block wild-type HSPCs. To enable long-term blockade of host HSPCs even after transplantation, we used prime editing to engineer CIM058-resistant human CD34^+^ HSPCs. When combined, CIM058 and the epitope engineered CD34^+^ HSPCs ameliorated disease phenotype in a β-thalassemia model. Our results suggest that this approach may overcome the reliance on busulfan or other myeloablative conditioning regimens with their associated morbidities, and by enabling toxin-free conditioning and in vivo selection of edited cells, may facilitate clinical implementation of these highly valuable genetic therapies.

## INTRODUCTION

Hematopoietic stem cell transplantation (HSCT) is a potentially curative approach for malignant and non-malignant diseases of the hematopoietic system^1,2^. Beyond conventional HSCT, ample clinical evidence demonstrates that ex vivo gene therapy and genome engineering of autologous hematopoietic stem and progenitor cells (HSPCs) can result in long-term functional correction of genetic defects underlying hematopoietic genetic diseases^3–7^. Importantly, for some indications such treatments could represent one-time cures which exemplify their immense potential^8^. Consequently, several genetically engineered HSPC products are now commercially available to treat diseases as diverse as metachromatic leukodystrophy, cerebral adrenoleukodystrophy, Wiskott-Aldrich syndrome, and the hemoglobinopathies sickle cell disease (SCD) and transfusion-dependent β-thalassemia (TDT)^9^.

β-Thalassemia is one of the most common inherited blood disorders worldwide, associated with substantial morbidity and early mortality, particularly in patients with limited access to supportive care or poor treatment adherence. Even in well-managed patients, quality of life is severely compromised by the life-long need for regular red blood cell transfusions and iron chelation. More than 300 mutations causing β-thalassemia have been identified, resulting in reduced or absent synthesis of the β-globin chains. The resulting imbalance between α- and β-globin chains leads to precipitation of excess α-globin, oxidative damage, and ineffective erythropoiesis characterized by impaired erythroid maturation, chronic anemia and compensatory expansion of erythropoiesis in the bone marrow and at extramedullary sites, resulting in splenomegaly. Chronic transfusions and dysregulated iron homeostasis cause progressive iron overload in vital organs, remaining a major cause of morbidity and mortality despite advances in supportive care^10–12^.

A deep understanding of globin gene expression regulation, developmental hemoglobin switching, and β-hemoglobinopathies pathophysiology has established β-thalassemia and sickle cell disease (SCD) as prototypical targets for curative genetic treatment approaches. Over the past decade, successive generations of versatile, precise and efficient genome engineering tools - including CRISPR/Cas9 nucleases, base editors (BE) and prime editors (PE) - were developed^13–18^ and rapidly validated therapeutically in clinical HSPC genome engineering trials for hemoglobinopathies^6,19,20^ and chronic granulomatous disease^21^. In fact, given the speed of the technical development and the impressive clinical success, it was recently proposed that correction of hemoglobinopathies has been reduced to an engineering problem^22^. However, while genome editing itself has become increasingly efficient, toxic conditioning remains a major barrier to broad clinical implementation of these otherwise curative therapies^23–26^.

A conditioning phase is required prior to the actual transplant to reduce or deplete the host HSPCs and create space for incoming donor HSPCs. However, current untargeted conditioning regimens for HSCT such as chemotherapy and/or irradiation, are directly or indirectly associated with substantial transplant-related morbidity and mortality^25^. Importantly, one of the conditioning molecules, the alkylating agent busulfan (Bu), was approved in the 1950’es. Its application has not changed much since then and it causes known severe toxicities including infertility, lung and liver toxicity and an increased risk for secondary malignancies^27,28^. The use of busulfan therefore constitutes a major safety concern, as illustrated in clinical trials of genome-edited HSPCs for hemoglobinopathies^19,20,29^ or a gene therapy for a primary immunodeficiency^8^, in which treatment-related deaths and serious adverse events were attributed to busulfan conditioning rather than the engineered cells themselves^8,19,20,29^. Furthermore, the risk for infertility can prevent young patients from opting for a potentially curative transplant^26^.

In summary, some of the most sophisticated and most recent engineered HSPC treatments rely on decades old, toxic conditioning which severely constrains safety, scalability and patient access. Therefore, there is a high medical need for alternative conditioning approaches to enable wide adoption of these curative therapies. Among the most promising strategies are antigen-specific immunotherapies targeting proteins expressed on HSPCs^30^. The receptor tyrosine kinase c-KIT (CD117) emerged as an attractive therapeutic target for conditioning prior to HSCT. It is activated by its ligand stem cell factor (SCF) leading to c-KIT signaling that is crucial for HSPC biology. Blocking SCF binding by the monoclonal antibody (mAb) SR-1 prevents proliferation in vitro^31^, while the anti-human CD117 mAb AMG191, a SR-1 derivative, was reported to induce partial and transient depletion of HSPCs in nonhuman primates (NHP) and human xeno-graft models in immunodeficient mice^32^. Thus, CD117-targeting, SCF blocking mAbs could represent a substantially less toxic alternative to conventional conditioning agents. A phase I clinical trial (NCT02963064) confirmed safety and a dose-dependent decline of CD117^+^ HSPCs but JSP191 (formerly AMG191) as a monotherapy for conditioning was insufficient to achieve substantial stable myeloid chimerism in patients with severe combined immunodeficiency (SCID) and a prior HSCT. Importantly, reduced myeloid chimerism was observed at the highest JSP191 dose^33^, indicating that a washout phase is needed before HSPC infusion to avoid depletion or prevention of engraftment of donor HSPCs by residual mAb^32,34,35^.

Alternatively, rather than waiting for the depleting mAb to fall below a threshold concentration^34,36^, one could render donor HSPCs resistant to the mAb. We and others recently demonstrated that epitope engineering of target proteins on HSPCs uncouples antigen-specific therapy from hematopoietic reconstitution after HSCT^37–40^. We, therefore, hypothesized that CD117 shielding on HSPCs would i) overcome the need for a mAb washout phase ii) possibly reduce time from conditioning to engineered HSPC infusion iii) enable mAb redosing post HSCT iv) allow extended in vivo selection and enrichment of CD117 shielded HSPCs and thus v) potentially create a titratable chimerism. Such an approach, could establish truly toxin-free conditioning, enabled by continuous SCF blockade of endogenous HSPCs.

Here, we identified CD117 variants that shield from a novel, concurrently developed, highly potent HSPC-depleting mAb (CIM058) while preserving normal CD117 function. Using prime editing, we introduced the lead variant CD117^E73K^ into human HSPCs. In a humanized β-thalassemia mouse model, conditioning with CIM058 followed by transplantation and engraftment of CD117^E73K^ HSPCs in combination with repeated post-transplant CIM058 administration, selectively enriched engineered cells in vivo, resulting in amelioration of β-thalassemia phenotypes.

## RESULTS

### Identification of CD117 variants that reduce SR-1 binding but preserve SCF binding

Alanine scanning of the CD117 (KIT) extracellular domain (ECD) was performed as described^37,41^ using an SR-1 Fab fragment to identify the epitopes targeted by the SR-1 antibody. SR-1 is the parent of briquilimab (formerly AMG191, later JSP191), a clinical-grade humanized antibody known to block stem cell factor (SCF) binding to CD117^31^. Two additional antibodies were used as antibodies of potential therapeutic interest and controls, namely Fab79D (Fab) and 104D2 (mAb) and YB5.B8 as technical expression control. Fab79D is a phage-derived antibody that binds to the membrane-proximal Ig-like domain D4 of CD117, preventing CD117 dimerization^42^. We noted that Fab79D aggregated when expressed in a human IgG1 format. Therefore, we only used Fab79D as control for preserved CD117 expression during Ala scanning but did not further pursue the antibody. 104D2 is a mouse monoclonal anti-CD117 antibody that has not been reported to inhibit SCF binding or CD117 dimerization. SR-1 epitope residues were identified as those where alanine substitution resulted in a maximum of 20% preserved SR-1 binding compared to the wild-type (WT) protein, while maintaining equal or greater than 70% binding for either of the two control antibodies (Figure 1A). This analysis identified the following SR-1 epitope residues: L56, L71, E73, G93, Y95, F110, D121, R122, S123, Y125, K127, V134, R181, S197, I201, K203 and R205, spanning protein domains D1-D3. As expected, Fab79D epitope residues are located on domain D4, while 104D2 epitope residues primarily cluster in a region of D1 distal and non-overlapping to the SR-1 epitope, with a few exceptions, e.g. E81A, Y95A, S197A. The latter, based on their protein and surface localization, are considered false positives. To further refine the SR-1 epitope, we prioritized residues where binding was largely preserved for all control antibodies (Figure 1A). As shown in Figure 1B and 1C, these residues lie within the SCF binding domain, with the majority being proximal to SCF interacting residues (Figure 1B and 1C). Residues at the periphery of the SCF binding region exhibit a less pronounced reduction in SR-1 binding upon alanine substitution, retaining more than 15% of their binding capability (e.g., L71, K127). An exception is E73, which shows approximately 5% residual binding to SR-1 upon mutation. E73 is located on a non-conserved loop, with its side chain pointing towards the protein surface, thus not involved in any key polar intra/intermolecular interactions. The latter was selected for further mutagenesis experiments (Figure 1C). Based on the observed effect of alanine substitutions and structural analysis of the CD117-SCF complex model, we selected several single amino acid substitutions for further characterization. Additionally, we arbitrarily included CD117 carrying a deletion of E73 (CD117^E73del^), CD117^E73P^ and the naturally occurring single nucleotide variant (SNV) CD117^S123F^.

**Figure 1:**
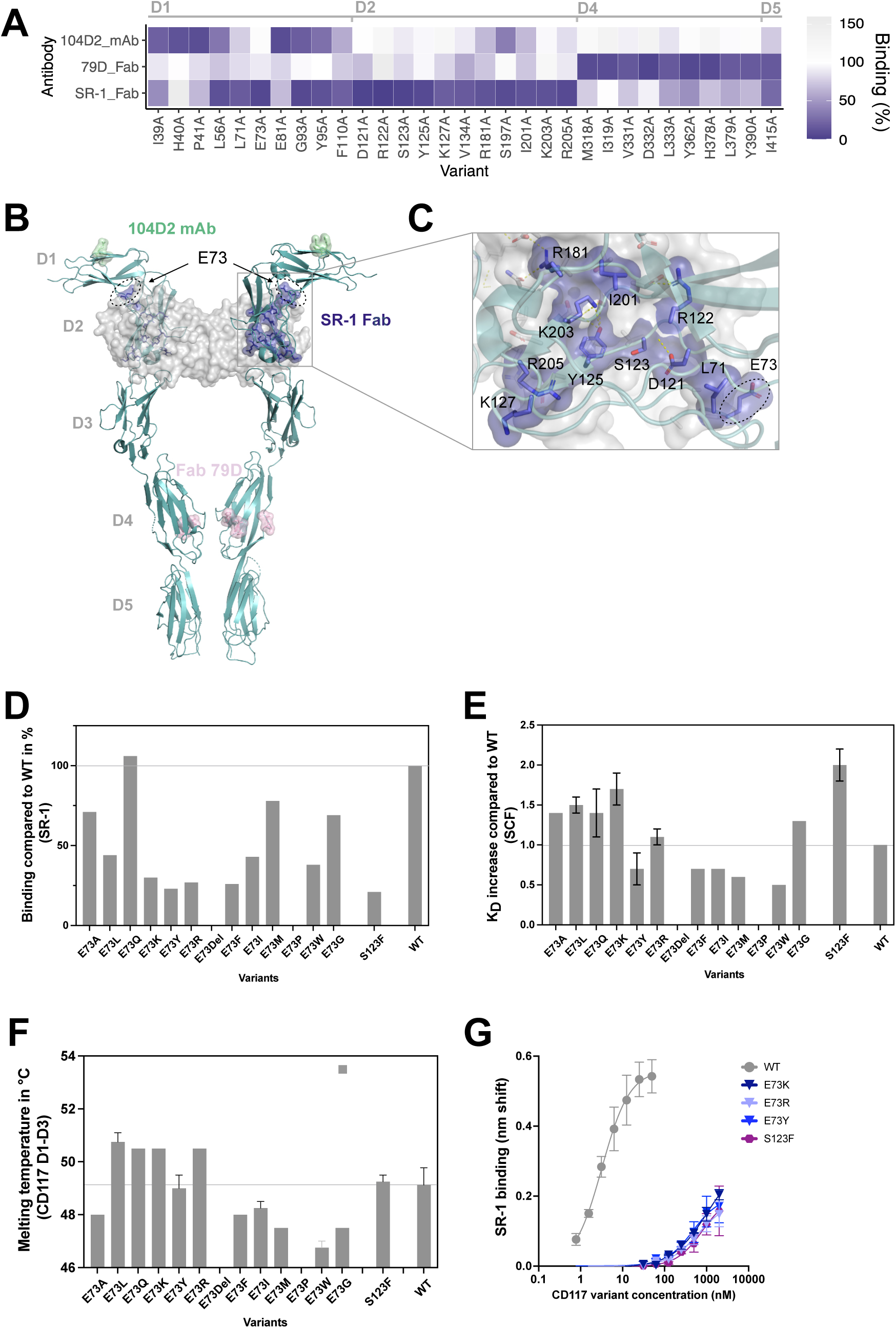
Identification of CD117 variants that reduce SR-1 binding but preserve SCF binding (A) Heatmap showing CD117 alanine substitutions that result in less than 20% residual binding for one monoclonal antibody but have more than 70% for either of the other two tested anti-CD117 antibodies. (B) 3D structure of the CD117-SCF complex (PDB ID: 2e9w). CD117 is shown as ribbons. SCF is shown as light grey molecular surface. CD117 residues involved in SCF binding are shown as dark grey surface. CD117 residues forming the SR-1 epitope are shown as purple surface. (C) Zoom on SR-1 epitope residues. Displayed as stick and surface. SCF shown as surface and residues in close proximity to CD117 (<5A) are shown as sticks. Polar contacts involving side chains displayed as dashed yellow lines. (D) Binding affinity of CD117 variants (ECDs D1-3) to SR-1 is shown as % binding vs WT. (E) Affinity (KD) of CD117 variants (ECDs D1-3) binding to SCF is shown as a ratio versus WT (biolayer interferometry). Values >1 (<1) indicate lower (higher) affinity to SCF. Error bars indicate the standard deviation of three measurements. Variant S123F is a naturally occuring SNV. (F) Melting temperature of purified recombinant CD117 protein (ECDs D1-3) variants. Square symbol corresponds to the 2^nd^ melting temperature of CD117^E73G^. (G) Dose titration of selected CD117 variants to test residual SR-1 binding at very high analyte concentrations. CD117 variants were chosen based on SR-1 binding reduction and preserved SCF binding. Note for panels D-E: CD117^E73del^ and CD117^E73P^ did not express and therefore there is no data point

To assess SR-1 binding to different CD117 variants, the domains D1-3 of the CD117 wild-type extracellular region as well as D1-3 containing selected mutations were produced as soluble, purified proteins. However, CD117 carrying a deletion of E73 (CD117^E73del^) and CD117^E73P^ did not express. To quantify the SR-1 binding reduction of selected CD117 variants compared to CD117^WT^, a label free real-time method was used. SR-1 antibody was captured on biosensors and the binding of CD117^WT^ or CD117 variants and their dissociation was monitored by biolayer interferometry (BLI). The percentage of binding compared to WT was calculated from the nm shift at the end of the association. Mutations that introduced positively charged (R, K) or bulky amino acids (F, Y, W, I, L) in position 73 reduced antibody binding the most, while the substitution to Q did not affect antibody binding (Figure 1D). This indicates that solely removing the negative charge (E to Q) does not suffice to reduce antibody binding. In contrast, inversion of charge from negative to positive or steric hindrance is sufficient. Furthermore, CD117^S123F^ also strongly decreased the degree of SR-1 binding (Figure 1D). An ideal shielding variant should reduce mAb binding but maintain its function. Therefore, binding of the selected variants to SCF was determined using BLI and compared to CD117^WT^. Biotinylated SCF was captured onto streptavidin coated biosensors. Similar to mAb binding, association and dissociation of CD117^WT^ and CD117 variants to immobilized SCF were measured. All mutations at position E73 as well as CD117^S123F^ retained SCF binding (Figure 1E). Since CD117^S123F^ represents a SNV, we assumed that the mild SCF binding variability observed for the E73 variants which is in a similar range to CD117^S123F^, may be biologically tolerated. To verify that the antibody shielding effect was a result of removing or disturbing a specific amino acid interaction rather than a result of major destabilization of CD117, CD117^WT^ and variants were analyzed by differential scanning fluorimetry (DSF). Most variants at position E73 displayed melting temperatures similar to CD117^WT^, suggesting that these substitutions did not markedly affect the overall thermal stability of the protein. We did not further pursue CD117^E73G^ because of a biphasic melting curve (Figure 1F).

To assess residual interactions, we measured SR-1 binding to CD117^WT^ and selected variants by BLI at very high analyte concentrations using a dilution series. Based on SR-1 binding reduction, preserved SCF binding and intact thermostability (Figures 1D-1F), we chose CD117^E73K^, CD117^E73R^, CD117^E73Y^ and CD117^S123F^. At a concentration where CD117^WT^ started to be saturated, the shielding variants did not display any SR-1 binding (Figure 1G). The titration curves illustrate a >1000-fold discrimination between CD117^WT^ and CD117^E73K^, CD117^E73R^, CD117^E73Y^ and CD117^S123F^ (Figure 1G). However, at high receptor concentrations, we detected some residual binding. In summary, we identified single amino acid substitutions (CD117^E73K^, CD117^E73R^, CD117^E73Y^ and CD117^S123F^) that strongly reduced binding to SR-1 even at very high concentrations but preserved CD117 protein stability and SCF binding.

### CD117 single amino acid substitutions retain SCF dependent function despite molecular shielding from SR-1 and derivatives

To investigate if CD117 variants affected mAb binding and to assess their potential biologic effects on cells, we used CRISPR/Cas9-based homology directed repair (HDR) to introduce CD117^E73K^ into the CD117 locus of the human SCF-dependent cell line TF-1. SR-1 binding determined by flow cytometry was abolished from two separate CD117^E73K^ expressing TF-1 clones whereas SR-1 showed the expected dose-dependent binding to unedited TF-1 cells (Figure 2A). In contrast, SCF bound in a dose-dependent manner to both unedited and CD117^E73K^ expressing TF-1 cells, respectively (Figure 2B). Edited TF-1 cells expressing CD117^E73K^ or CD117^S123F^ retained SCF-dependent growth with EC50 values comparable to WT TF-1 cells (Figure 2C). SR-1 dose-dependently inhibited cell growth of unedited TF-1 cells while engineered TF-1 expressing CD117^E73K^ or CD117^S123F^ were unaffected by the presence of SR-1 (Figure 2D). Next, we measured SCF induced signal transduction by quantifying Y719 phosphorylation (p-CD117^Y719^). First, we performed a SCF dose titration (Figure S1A). Wild-type TF-1 displayed a clear dose-dependent CD117^Y719^ phosphorylation, while removing CD117 (CD117^KO^) abolished SCF signaling entirely. In contrast, CD117^E73K^ clones 1 and 2 both displayed dose-dependent CD117^Y719^ phosphorylation (Figure S1A), indicating normal SCF sensing and signal transduction. Next, we tested SCF induced p-CD117^Y719^ in the presence of SR-1. Wild-type TF-1 cells were responsive to SCF; SR-1 alone did not induce any p-CD117^Y719^ but SR-1 efficiently blocked SCF from inducing p-CD117^Y719^ (Figure 2E). CD117^KO^ were unresponsive to SCF, SR-1 or the combination of both. In contrast, SR-1 was unable to block SCF-induced p-CD117^Y719^ on cells expressing CD117^E73K^ in both clones tested (Figure 2E)

**Figure 2:**
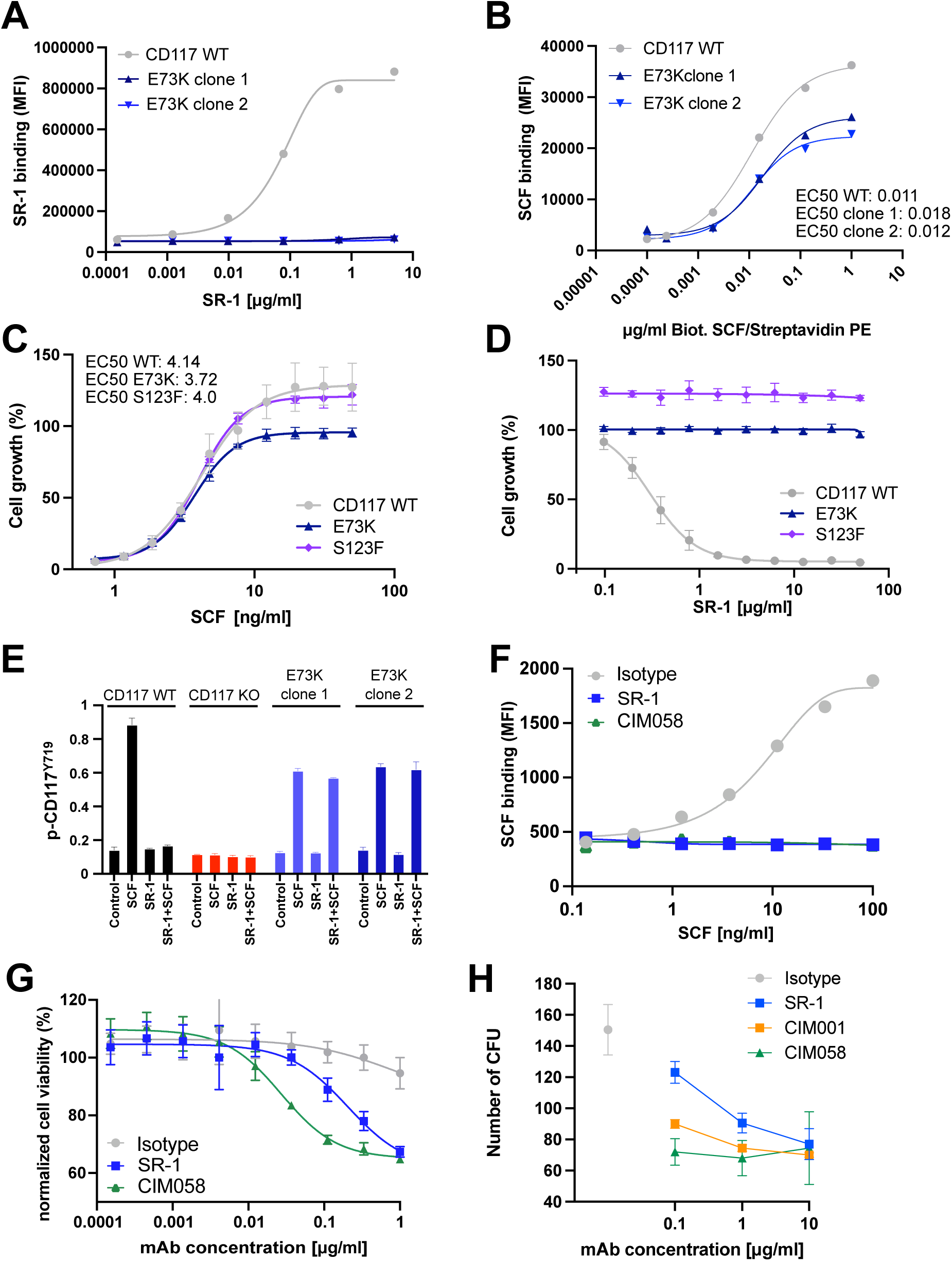
CD117 single amino acid substitutions retain SCF dependent function despite molecular shielding from SR-1 and derivatives (A) Binding of the anti-CD117 antibody SR-1 to TF-1 cells expressing engineered CD117 variant E73K in comparison to cells expressing the wildtype receptor. Shown are results from two independent TF-1 E73K clones. (B) Binding of the SCF to TF-1 cells expressing engineered CD117 variant E73K in comparison to cells expressing the wildtype receptor (EC50). Shown are results from two independent TF-1 E73K clones. (C) SCF dose-dependent proliferation of TF-1 cells expressing engineered CD117 variant E73K and the natural SNV S123F in comparison to cells expressing the wildtype receptor (EC50). (D) SCF-mediated growth in the presence of the anti-CD117 antibody SR-1 of TF-1 cells expressing wildtype or the engineered variants E73K and S123F. (E) CD117 phosphorylation at position Tyr719 upon anti-CD117 antibody treatment in TF-1 cells expressing wildtype, KO CD117 or the engineered variant E73K; SCF: 0.1µg/ml; SR-1: 0.5µg/ml. n = 1 biological replicate with 2 technical replicates (F) Binding of SCF to HSPCs in the presence of anti-CD117 antibodies (50µg/ml). (G) Proliferation of mobilized CD34^+^ HSPCs in the presence of anti-CD117 antibodies. (H) HSPCs were plated on methylcellulose with increasing concentrations of anti-CD117 antibodies or 10 μg/ml isotype control and allowed to grow for 2 weeks in the presence of anti-CD117 antibodies. Total number of colonies were counted and shown as mean ±SD (n=1 experiment, 2 replicates). See also Figure S1.

The SR-1 derivative briquilimab has shown limited efficacy as a standalone conditioning agent for HSPC depletion in HSCT clinical trials^35^. Therefore, we aimed to generate improved, more potent SCF blocking mAbs. In a first step we humanized the SR-1 variable regions and incorporated them into an Fc silent human IgG1 (CIM001) because co-engagement of CD117 and FcγR was described to be responsible for mast cell degranulation which occurred in a clinical trial using a CD117-targeting antibody with an unmodified Fc^43^. CIM001 bound CD117^WT^ similar to SR-1 and also displayed reduced binding to the selected variants CD117^E73K^, CD117^E73R^, CD117^E73Y^ and CD117^S123F^ (Figure 1G and S1B, left panel). Next, we increased mAb affinity and removed liabilities such as multiple sites that are prone to post-translational modifications or fragmentation, resulting in CIM058. This improved mAb still displayed strongly reduced binding to the selected variants (Figure S1B, right panel). Next, we confirmed that SCF-dependent phosphorylation of CD117 remained intact in TF-1 cells expressing CD117^E73K^ in the presence of SCF and CIM001 (Figure S1C). In contrast, CIM001 abolished SCF-dependent phosphorylation in CD117^WT^ TF-1 cells. Finally, we confirmed that CIM001 and CIM058 abolished SCF binding to CD117^WT^ TF-1 cells, whereas isotype exposure did not hinder SCF binding. In contrast, TF-1 cells expressing CD117^E73K^ equally bound SCF in the presence of isotype, SR-1, CIM001 or CIM058 (Figure S1D).

Importantly, we tested SCF binding to primary human HSPCs in the presence of CD117 antibodies or isotype control (Ref001). SCF bound in a dose-dependent manner in the presence of the isotype control. The addition of SR-1 or CIM058 completely abolished SCF binding, confirming the results obtained with TF-1 cells (Figure 2F). Furthermore, HSPC growth was inhibited in a dose-dependent manner by SR-1, and the inhibition was even more pronounced with the high-affinity antibody CIM058 (Figure 2G). Finally, HSPC colony formation in methylcellulose was reduced in a dose-dependent manner upon addition of the antibodies, with the effect of SR-1 being less pronounced than that of CIM001 and CIM058, demonstrating increased antibody potency (Figure 2H). These results demonstrated that CD117^E73K^ bound SCF and signalled normally, but was shielded from binding to the antibodies SR-1, CIM001 and CIM058, respectively.

### The CD117 antibody CIM058 enriches HSPCs expressing CD117^E73K^ in vitro

With clinical translation in mind, we next engineered CD117^E73K^ into clinically relevant CD34^+^ HSPCs collected by apheresis. HSPCs were electroporated with high-fidelity SpCas9 ribonucleoproteins (RNPs) and single-stranded oligodeoxynucleotide (ssODNs) donor templates, using an editing strategy similar to the one we recently described for CD123 epitope engineering^38^. CD117 expression and editing were monitored using flow cytometry with the mAbs 104D2 (control antibody clone) and SR-1. In electroporation only control HSPCs (“EP” hereafter), most cells were 104D2^+^ SR-1^+^ double positive (DP). We did not detect 104D2^+^ SR-1^-^ single positive (SP) cells but some 104D2^-^ SR-1^-^ double negative (DN) cells (Figure 3A). In contrast, SP cells were present in HSPCs electroporated with the RNP plus CD117^E73K^ ssODN template (“E73K” hereafter). We also noted that the DN fraction increased in E73K cells (Figure 3A). Compared to bulk CD34^+^ cells, frequencies of SP cells were slightly reduced in phenotypic LT-HSCs (CD34^+^CD38^-^CD90^+^CD45RA^-^) (Figure 3B). Next, we analyzed editing outcomes by next-generation sequencing (NGS). Control HSPCs (EP) were nearly completely wild-type (Figure 3C), indicating that the DN cells in the EP group (Figure 3A) were HSPCs that did not express CD117 but had an intact CD117 locus. In contrast, CD117^E73K^ knock-in (KI) reads were confirmed in E73K cells, and the editing efficiency (KI) correlated well with flow cytometry data (SP). Besides KI, we detected WT reads and indels (KO), suggesting that the increased DN fraction represented true KO cells (Figures 3A and 3C). To investigate whether the edited HSPCs were functional and could be enriched in vitro upon antibody treatment, a colony forming assay was performed using EP or E73K HSPCs in the presence of CIM058 or the isotype control antibody Ref001. E73K HSPCs formed a similar number of colonies as EP HSPCs in the presence of Ref001 (Figure 3D). Adding CIM058 reduced the number of colonies, an effect which was more pronounced in the EP control group. To quantify the allele frequency and verify if CIM058 enriched KI alleles, colonies were picked and subjected to Sanger sequencing (Figure 3E). Colonies from the EP group displayed only WT reads, regardless of whether Ref001 or CIM058 was present in the culture. Conversely, different alleles were found in colonies derived from E73K-edited cells: WT/WT, WT/KI, KI/KI, KI/KO, WT/KO and KO/KO. Compared to Ref001 treated cells, CIM058 clearly decreased the number of WT/WT and WT/KO colonies but enriched genotypes harbouring the E73K allele, namely KI/KI and KI/KO. These results demonstrated that cells expressing CD117^WT^ were depleted whereas CD117^E73K^ expressing cells were enriched by CIM058 treatment in vitro. To assess the safety of HDR-mediated CD117^E73K^ engineering, and nominate potential off-target (OT) sites of the nuclease used, we performed CAST-Seq^44^ (Figure 3F). Irrespective of the presence of ssODN, CAST-Seq did not detect any OT-mediated translocations (OMTs) that would indicate the existence of off-target activity. However, under both conditions, large deletions and inversions affecting a region of approximately 10 kb around the cut site were detected, which are absent in the non-electroporated control sample (No EP). In conclusion, the CD117^E73K^ allele - when engineered into HSPCs - conveyed resistance to CIM058, which enriched CD117^E73K^ engineered cells among the pool of electroporated HSPCs. Furthermore, initial assessment demonstrated a favorable safety profile.

**Figure 3:**
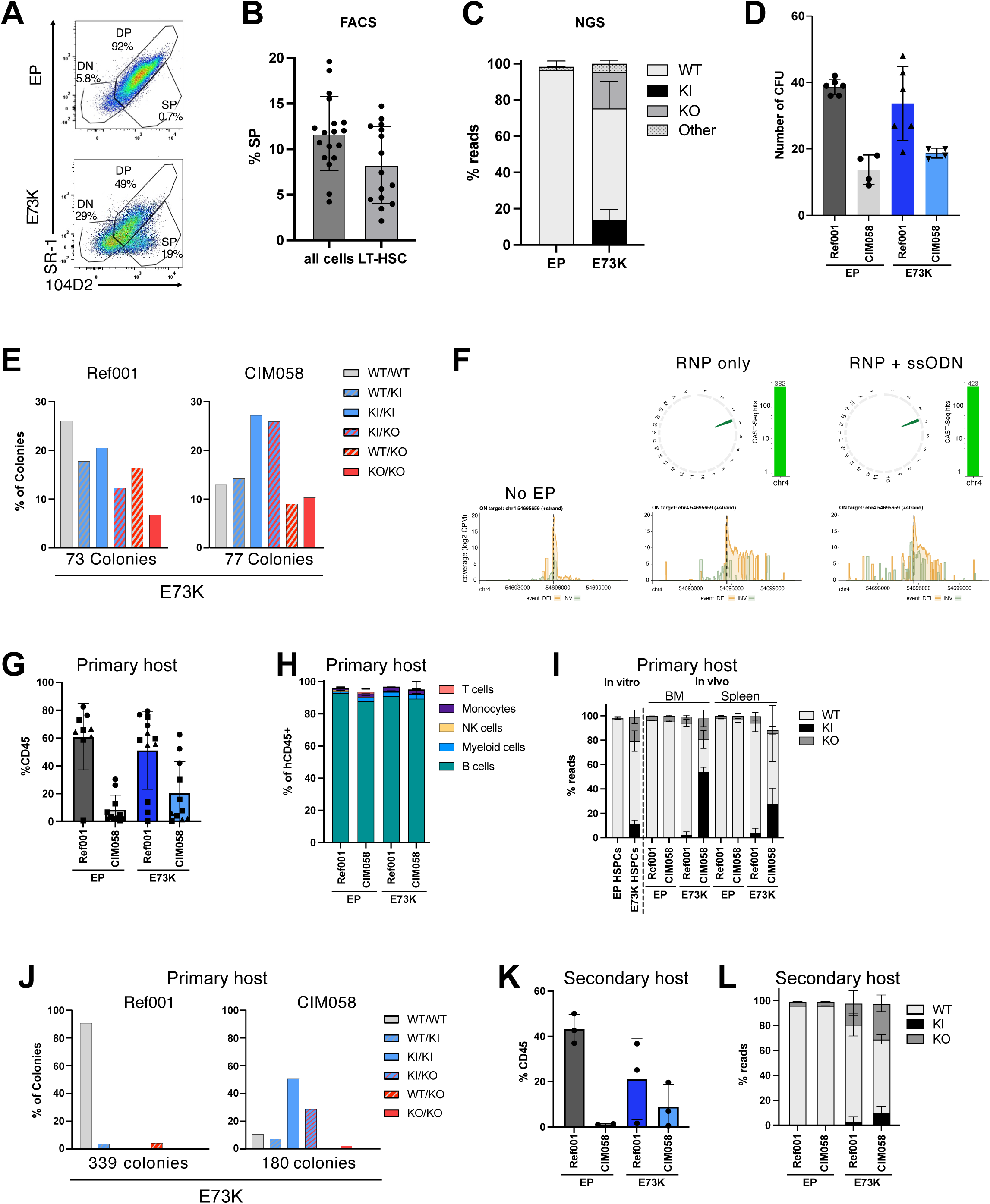
CIM058 enriches HSPCs expressing CD117^E73K^ in vitro and CD117-shielded HSPCs engraft and differentiate normally (A) Characterization of HSPC variant E73K engineered with CRISPR/Cas9 HDR by FACS using the control antibody 104D2 and SR-1. (B) Quantification of the edited cells based on the FACS gated on all viable cells or LT-HSC. (C) Quantification of the alleles by NGS. KI=E73K; Other=indels (D) Control electroporated HSPCs or E73K edited HSPCs were plated on methylcellulose and allowed to grow for 2 weeks in the presence of Ref001 control or CIM058 anti-CD117 antibody. Total number of colonies were counted. (E) Colonies from (D) were picked and subjected to Sanger sequencing. (F) Circos plots and coverage plots of CAST-seq results of no EP, RNP only and RNP+ssODN: ON aberrations are shown in green. (G-I) NBSGW mice were engrafted with EP control or E73K edited HSPCs. One week after HSPC injection (D7, D9, D11 and D11), mice received 4 doses of Ref001 or CIM058 (4mg/kg). 16 weeks later mice were euthanised and blood, spleen and BM were analysed. Flow cytometry analysis of human CD45^+^ in the BM (G), multi-lineage differentiation in the spleen (H) and quantification of the WT, KO and edited reads in the HSPCs before in vivo injection and in the spleen and BM of the mice (I). Shown are results from 3 different experiments performed with HSPCs from different donors (different symbol for each donor). (J) BM cells from primary transplant were plated on methylcellulose and allowed to grow for 2 weeks. Colonies were picked and subjected to Sanger sequencing. (K-L) Secondary trasplant experiment. 40% of BM from the primary trasplant was injected in irradiated NSG-SGM3 mice. Eight weeks later mice were euthanised and spleen and BM were analysed. Flow cytometry analysis of human CD45^+^ in the BM (K) and quantification of the WT, KO and edited reads in the spleen of the mice (L).

### CD117-shielded HSPCs engraft and differentiate normally, demonstrating long-term reconstitution potential in vivo

To test the in vivo engraftment and differentiation potential of CD117-engineered HSPCs, we injected EP and E73K-edited cells separately into immunodeficient NBSGW mice. One week after injection, the mice received 4 injections of either Ref001 or CIM058. After 16 weeks, comparable engraftment of hCD45^+^ cells was observed in the bone marrow of mice engrafted with either EP control or E73K cells treated with Ref001 (Figure 3G). We observed some engraftment variability, which was more pronounced in one HSPC donor (squares) compared to the two others (circles and triangles). Variability may be related to slightly different electroporation conditions and impaired cell viability caused by HDR-mediated HSPC engineering, as previously reported^38^. Importantly, treatment with CIM058 reduced the frequency of hCD45^+^ cells for both conditions, with a greater reduction observed in mice that received EP HSPCs compared to those engrafted with E73K HSPCs (Figure 3G). All groups exhibited similar multi-lineage differentiation in the spleen (Figure 3H). To quantify the presence and possible enrichment of CD117^E73K^ alleles, we performed NGS on BM and spleen. Prior to injection, cells displayed 11% KI reads and nearly 20% KO reads in the E73K-edited HSPCs (Figure 3I). Mice receiving EP cells did not show KI reads, independent of organ and treatment group. In mice engrafted with E73K cells and treated with control Ref001, the percentage of E73K edited reads in the BM and spleen dropped to a few percent while the percentage of KO reads remained stable. In contrast, CIM058 treatment strongly enriched the KI reads in BM and spleen, reaching >50% in BM. WT reads decreased but compared to Ref001, the KO reads were also enriched, especially in the BM. Colony forming assays using BM cells from primary recipients largely confirmed these results. Colonies derived from BM of Ref001 treated mice were nearly exclusively WT/WT with a few WT/KI and WT/KO genotypes. In contrast, the genotype of colonies derived from CIM058 treated mice was predominantly KI/KI and the second most frequent genotype was KI/KO. WT/WT, WT/KI, WT/KO and KO/KO were rather rare (Figure 3J).

Finally, secondary transplantation experiments from one donor showed engraftment of human cells in BM, blood and spleen. Mice engrafted with EP control cells and treated with Ref001 showed around 40% hCD45 cells in BM whereas CIM058 treatment of the primary EP cell recipients prevented engraftment in secondary hosts nearly completely (Figure 3K). This demonstrated that CIM058 effectively depleted LT-HSCs in primary host mice. Secondary engraftment of mice receiving E73K HSPCs was lower compared to EP but hCD45^+^ cells were present in 2/3 secondary host mice in the E73K group despite CIM058 treatment (Figure 3K). NGS neither showed KO nor KI reads in the EP groups whereas KI reads were detected in the E73K groups, with more reads after CIM058 treatment (Figure 3L). However, compared to the primary hosts (Figure 3I), the relative number of KI reads decreased while KO reads remained constant or even increased (Figure 3L).

In conclusion, HDR-mediated CD117^E73K^ HSPC engineering was feasible but viability and long-term engraftment potential were impaired which could be related to the variant itself or the editing approach. Furthermore, KO cells were generated that persisted unexpectedly long-term. Nevertheless, the results clearly demonstrated that CD117^E73K^ HSPCs can persist long-term and CIM058 selectively depletes WT cells resulting in long-term in vivo enrichment of cells harboring a CD117^E73K^ genotype.

### Prime editing improves CD117^E73K^ epitope engineering in HSPCs

To overcome the limitations of CRISPR/Cas9 nuclease-induced HDR (e.g. impaired viability, long-term engraftment capacity and heterogenous editing outcomes including KI and KO cells), we tested BE, which was recently shown to be effective for epitope engineering and result in high cellular viability^37,39,40^. However, preliminary experiments using two cytosine base editors (CBE4max-SpRY^45^ and TadCBEa(V106W)-SpRY)^46^ generated low on-target CD117^E73K^ editing efficiency and similar unintended bystander editing resulting in additional amino acid substitutions (Figure S2A). Since bystander editing might impair SCF binding, we set up PE as a templated approach that can avoid indels and bystander edits^16^ and is well tolerated if delivered as RNA (high viability). A first round of epegRNA design optimization resulted in “epeg4”, containing the desired edit for the E73K substitution as well as a silent PAM-disrupting mutation (Figure 4A). Additionally, we designed a nicking guideRNA (ngRNA) on the reverse strand that can only cut once the PAM mutation and the CD117^E73K^ codon have been installed. Control cells were generated using an epegRNA encoding a PAM mutation and a silent mutation at position E73 (“epegControl”). HSPCs that had been electroporated with the epegRNA, the ngRNA and PEmax^47^ were used for in vitro differentiation and CFU assays. The edited cells were grown in high cytokine media with or without CIM058, and analysed by NGS after 2 weeks. CIM058 treatment depleted CD117^epegC-PE^ HSPCs whereas the CD117^E73K-epeg4-PE^ allele was enriched twofold (Figure 4B). Importantly, we did not detect any indels or bystander edits in either the saline group or the CIM058-enriched cells, which is in contrast to HSPCs engineered by HDR or BE. The CD117^epegC-PE^ and CD117^E73K-epeg4-PE^ HSPCs formed a comparable number of colonies, and the erythroid-to-myeloid ratio was similar (Figure 4C). Next, we tested in vivo engraftment and enrichment potential, using CD117^epegC-PE^ and CD117^E73K-epeg4-PE^ HSPCs injected into NBSGW mice (Figure S2B). To investigate the potential for enrichment of low frequencies of engineered cells, we used CD117^E73K-epeg4-PE^ HSPCs with a starting editing rate <1% (Figure S2C). The Mice received a total of three CIM058 or saline injections (3 x 4mg/kg), and enrichment was assessed by NGS in blood samples collected eight weeks after HSPCs injection. In the mice engrafted with CD117^epegC-PE^ cells, treatment with CIM058 depleted control edited alleles, whereas the edited alleles were slightly enriched in mice injected with CD117^E73K-epeg4-PE^ cells (Figure S2C). At the end of the experiment, engraftment of CD117^epegC-PE^ and CD117^E73K-epeg4-PE^ cells was comparable in BM and spleen in the saline group (Figure S2D). Treatment with CIM058 resulted in a reduction of hCD45^+^ cells, which was more pronounced in mice receiving the CD117^epegC-PE^ cells than the CD117^E73K-epeg4-PE^ cells. NGS analysis of BM and spleen confirmed enrichment of CD117^E73K-epeg4-PE^ cells upon CIM058 treatment from the starting 1% edited reads to nearly 5% in the BM and 13% in the spleen, demonstrating effective enrichment even when the initial editing is low (Figure S2E). Encouraged by these results, we aimed to further increase the editing efficiency. To this end, we introduced an additional silent mutation between the PAM and E73 codon in epeg4, thus generating epeg14, and adapted the ngRNA sequence accordingly (Figure 4D)^47^. As, assessed by flow cytometry, epeg14 increased the editing efficiency (Figure 4E). Of note, in contrast to HDR (Figure 3A), PE-mediated CD117^E73K-epeg14-PE^ engineering did not increase the frequency of DN cells compared to the EP, indicating that few or no indels were generated. In an in vitro differentiation assay, NGS confirmed the absence of indels despite increased editing efficiency of epeg14 compared to epeg4 in the saline groups (Figure 4F). Moreover, CIM058 enriched cells engineered with epeg4 or epeg14 but not epeg control (Figure 4F). Lastly, for an initial unbiased analysis of indels and chromosomal translocations, we used CAST-Seq to enable a direct comparison to the results obtained with HDR (Figure 3F). Again, CAST-Seq did not detect putative OT effects. Furthermore, and in contrast to the nuclease approach, no evidence of gross chromosomal rearrangements at the on-target site in CD117^E73K-epeg14-PE^ samples were observed (Figure 4G). In conclusion, prime editing was equally or more efficient than HDR for CD117^E73K^ engineering into HSPCs and did not induce unintended indels or bystander edits, resulting in a highly precise editing outcome. CD117^E73K^ prime edited cells differentiated like control cells in vitro and provided equal shielding and enrichment potential with CIM058 treatment in vitro and in vivo.

**Figure 4:**
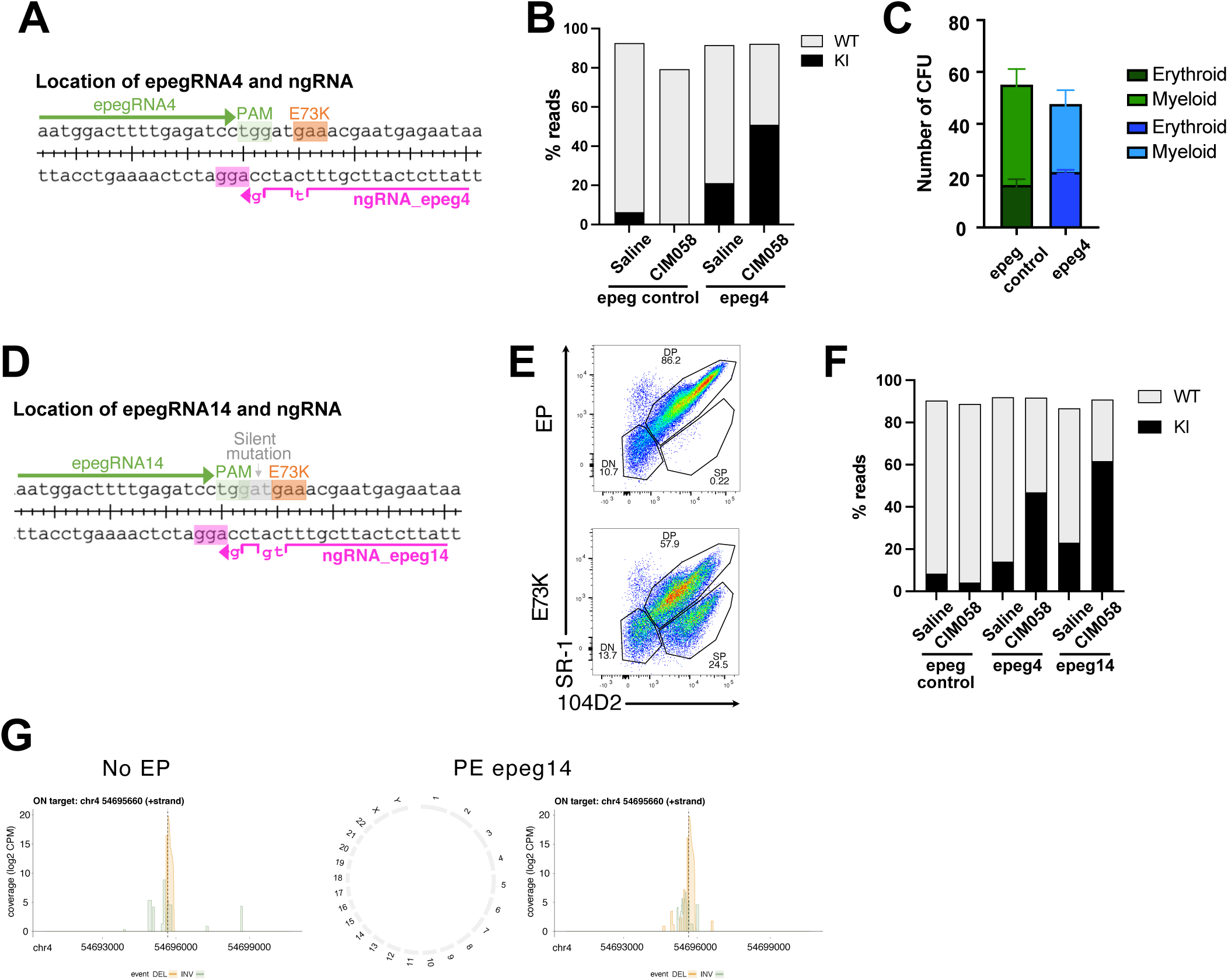
Prime editing improves CD117^E73K^ epitope engineering in HSPCs (A) Schematic representation of the position of the epegRNA4 and nicking guide. (B) In vitro differentiation of control-edited cells (epeg control introduces silent and PAM mutations) and E73K-edited cells (epegRNA4 introduces the E73K mutation and a PAM mutation) was examined in the presence of saline or the CIM058 antibody. The cells were subjected to NGS analysis two weeks after culture. (C) Control electroporated HSPCs or E73K edited HSPCs were plated in methylcellulose and grown for two weeks. The total number of myeloid and erythroid colonies was counted. (D) Schematic representation of the position of the epegRNA14 and nicking guide. (E) Characterization of HSPC variant E73K engineered with epeg14 and ngRNA by FACS using the control antibody 104D2 and SR-1. (F) In vitro differentiation of control-edited cells (epeg control introduces silent and PAM mutations) and epeg4 E73K-edited cells or epeg14 E73K-edited cells (epegRNA14 introduces the E73K mutation, a silent mutation and a PAM mutation) was examined in the presence of saline or the CIM058 antibody. The cells were subjected to NGS analysis two weeks after culture. (G) Circos plots and coverage plots of CAST-seq results of no EP control cells and E73K epegRNA14 edited cells. See also Figure S2.

### CD117 shielded HSPCs retained function and could be enriched in vivo

Next, we focused on in vivo behaviour of prime edited HSPCs. We engrafted CD117^epegC-PE^ or CD117^E73K-epeg14-PE^ HPSCs to NBSGW mice and divided them after 11 days into 4 different treatment groups: PBS treatment (group 1) or 3 different CIM058 treatment regimens (groups 2-4). Group 2 received 4 x 4mg/kg early doses of CIM058 on days d0, d2, d4 and d6, starting 11 days after HSPC engraftment (=d0) (Figure 5A). Group 3 received 4 x 1mg/kg on d0, d14, d28 and d42, i.e. was also started early but follow-up doses were bi-weekly whereas group 4 received the first CIM058 dose on d28 and then an additional 3 x 1mg/kg bi-weekly injections (Figure 5A). In contrast to the pan-hematopoietic marker CD45 which allows straightforward monitoring of epitope engineering in peripheral blood by FACS^37^, CD117 is only expressed on rare cells, making it challenging to quantify CD117 epitope engineering by flow cytometry from limited volumes of murine blood. Therefore, we performed NGS on peripheral blood samples. In mice treated with PBS, CD117^E73K^ remained stable from d27-d94, demonstrating that CD117^E73K^ neither conveyed a competitive advantage nor disadvantage, i.e. was functionally neutral compared to control CD117^epegC-PE^ HSPCs (Figure 5B). In contrast, mice that received CIM058 injections demonstrated increasing CD117^E73K^ reads over time, indicating a clear selection by CIM058. Of note, since only few cells express CD117 in peripheral blood, this increase detected by NGS most likely reflected selection on a precursor level in the BM which resulted in delayed enriched blood cells carrying the engineered genome, even when they do not express CD117. In line with this interpretation, the mice that received late CIM058 injections (group 4) demonstrated delayed and lower CD117^E73K^ reads in peripheral blood (Figure 5B). Mice receiving 4 x 4 mg/kg (group 2) or 4 x 1 mg/kg (group 3) displayed similar kinetics, showing around 40% edited cells by day 49 (i.e. a doubling of CD117^E73K^ reads), and almost 60% by day 94. This suggests that a higher dose was not required to achieve the same effect. Mice receiving late dosing showed a near-doubling of CD117^E73K^ reads after 66 days, suggesting that the effect in this group is probably only delayed, but not different.

**Figure 5:**
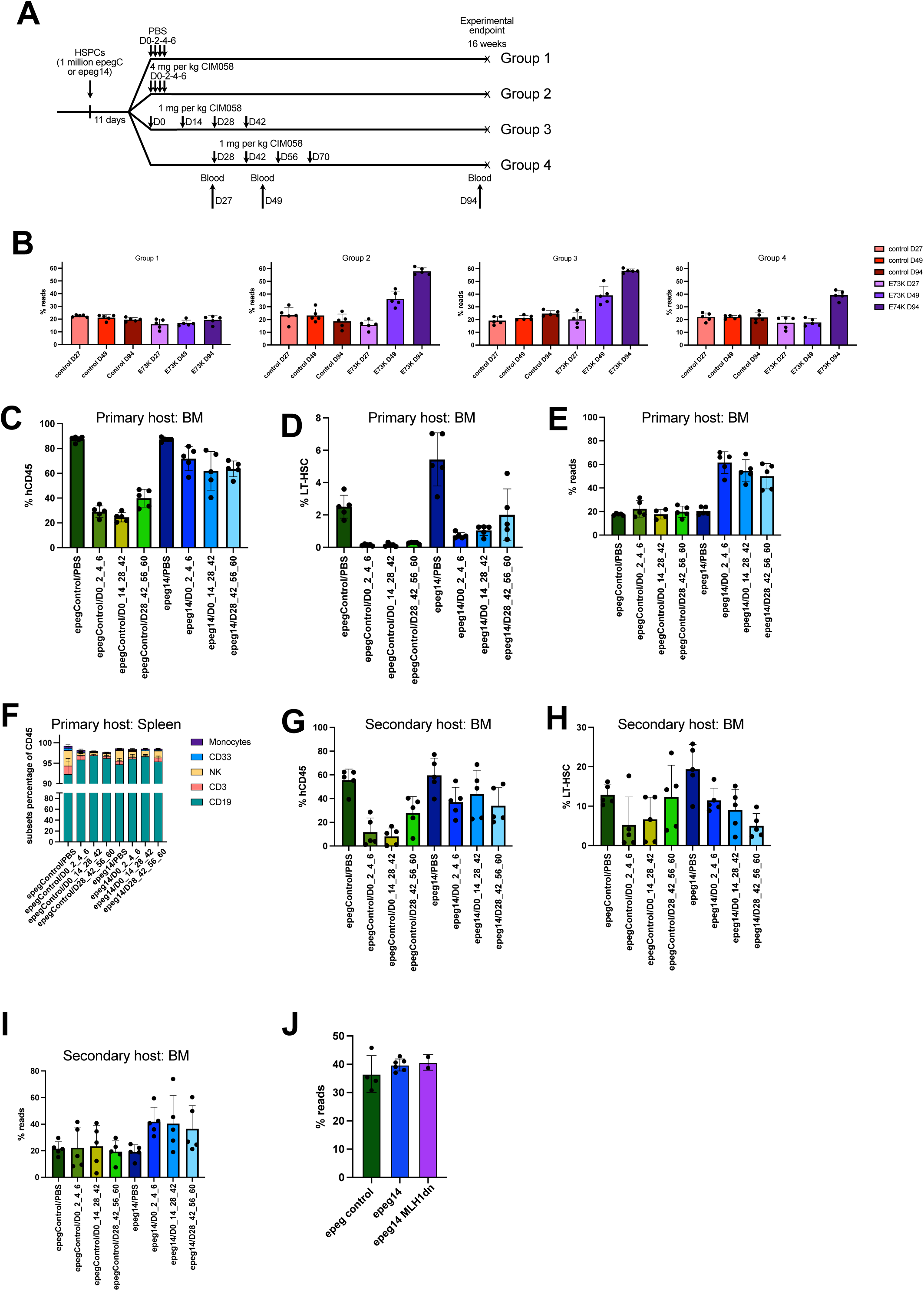
CD117 shielded HSPCs retained function and can be enriched demonstrating long-term reconstitution potential in vivo (A) Schematic representation of the mouse in vivo experiment. NBSGW mice were engrafted with epegRNA control or E73K epegRNA14 edited HSPCs and euthanised after 16 weeks. Eleven days after cell injections mice received 4 injections every 2 days of PBS (Group 1) or CIM058 4mg/kg (Group 2) or every 2 weeks of CIM058 1mg/kg (Group 3). Group 4 received 4 injections every 2 weeks of CIM058 1mg/kg starting at day 28 after HSPC injection. Human chimerism was analysed in the blood 27, 49 and 94 days after HSPC injection. (n = 5 mice per group and HSPC genotype) (B) NGS analysis of the edited reads (KI for control: silent and PAM mutations; KI for E73K: E73K codon, silent and PAM mutations) in the blood of the mice at day 27, 49 and 94. (C-D) FACS quantification of human CD45^+^ cells (C) or LT-HSC in the BM of the mice at the end of the experiment. (E) NGS analysis of the edited reads in the BM of the mice at the end of the experiment. (F) Flow cytometry analysis of the multi-lineage differentiation in the spleen of the mice at the end of the experiment. (G-I) Secondary transplant experiment. 40% of BM from the primary transplant was injected in irradiated NSG-SGM3 mice. Eight weeks later mice were euthanised and spleen and BM were analysed. Flow cytometry analysis of human CD45^+^ (G) and LT-HSC (H) in the BM of the mice. Quantification of the edited reads (I) in the spleen of the mice at the end of the experiment. (J) NGS analysis of the edited reads in HSPCs electroporated with epeg control, epeg14 or epeg14 together with MLH1dn. n = 2-6 biological replicate performed with 3 different donors See also Figure S3.

Analysis of the BM, spleen and blood revealed similar levels of engraftment of the CD117^epegC-PE^ and CD117^E73K-epeg14-PE^ HSPC in group 1 (Figure 5C and S3A and S3B). Treatment with different CIM058 regimens resulted in a substantial reduction in the number of hCD45^+^ cells in mice humanised with CD117^epegC-PE^. In contrast, mice engrafted with CD117^E73K-epeg14-PE^ HSPCs showed minimal cell depletion, indicating successful shielding of the edited cells (Figure 5C). Importantly, the LT-HSCs of mice humanised with CD117^E73K-epeg14-PE^ cells were also protected from CIM058 treatment (Figure 5D). Relative editing frequencies evaluated by NGS remained constant in the BM and the spleen of CD117^epegC-PE^ mice, whereas CIM058 injection reduced CD117^WT^ reads, resulting in an enrichment of E73K-edited cells (Figure 5E and S3C). Cell type analysis in the spleen and blood revealed similar multi-lineage cell differentiation potential across all groups and treatments (Figures 5F and S3D). Secondary transplantation experiments showed high and equal BM engraftment of CD117^epegC-PE^ and CD117^E73K-epeg14-PE^ cells in group 1 (Figure 5G) and clearly detectable LT-HSCs (Figure 5H) eight weeks after HPSC injection. CIM058-treated CD117^E73K-epeg14-PE^ cells engrafted more similarly to the PBS control group while CIM058-treated CD117^epegC-PE^ ^cells^ exhibited reduced engraftment (Figures 5G and 5H), reflecting the depletion of the unedited LT-HSC in the primary host mice (Figure 5D). Human cells were also present in the spleen (Figure S3E). Importantly, CIM058 administration in the primary host did not impair secondary engraftment of shielded HSPCs and all mice engrafted (Figures 5G and 5H). NGS analysis confirmed that relative editing frequencies remained constant in the spleen of the secondary hosts, confirming the persistance and functionality of the CD117^E73K-epeg14-PE^ HSPCs (Figure 5I). Compared to the secondary transplantation results with the HDR-engineered HSPCs (Figure 3K), these results underscored that the true long-term repopulation potential and LT-HSC function was more preserved in prime edited HSPCs. Given these in vivo data, we revisited alternatives to increase PE efficiency, including the use of a dominant negative MLH1 (MLH1dn)^47^. Although MLH1dn did not increase the CD117^E73K-epeg14-PE^ efficiency, we noted higher editing than in previous experiments (Figure 5L). This illustrates the possibility to further increase the editing efficiency.

### Shielded HSPCs paired with CIM058 ameliorated disease phenotype in a humanized β-thalassemia mouse model

Finally, we investigated the therapeutic potential of non-toxic antibody conditioning combined with in vivo selection in a disease model of allogeneic HSCT. We tested whether CIM058 could deplete patient-derived β-thalassemia CD34^+^ HSPCs and subsequently enrich transplanted, CD117-shielded HSPCs carrying a WT β-globin gene. To establish the disease, NBSGW mice were engrafted with CD34^+^ cells from a β-thalassemia patient with a homozygous IVSI-110/IVSI-110 mutation^48^ (hereafter “β-thal”), while mice engrafted with healthy donor (HD) HSPCs served as controls. Following characterization of the disease phenotype at baseline, β-thal recipient mice were split to receive either two doses of Ref001 isotype control, or two doses of CIM058 as antibody-based conditioning. Two weeks later, both groups were transplanted with CD117^E73K-epeg14-PE^ HSPCs from an allogeneic healthy donor (HSCT), followed by repeated Ref001 or CIM058 injections. The treatment effect was monitored by serial blood sampling before terminal analysis 9 weeks after the first post-transplant antibody injection. The overall experimental setup consisted of 8 experimental groups including all controls (Figure 6A).

**Figure 6:**
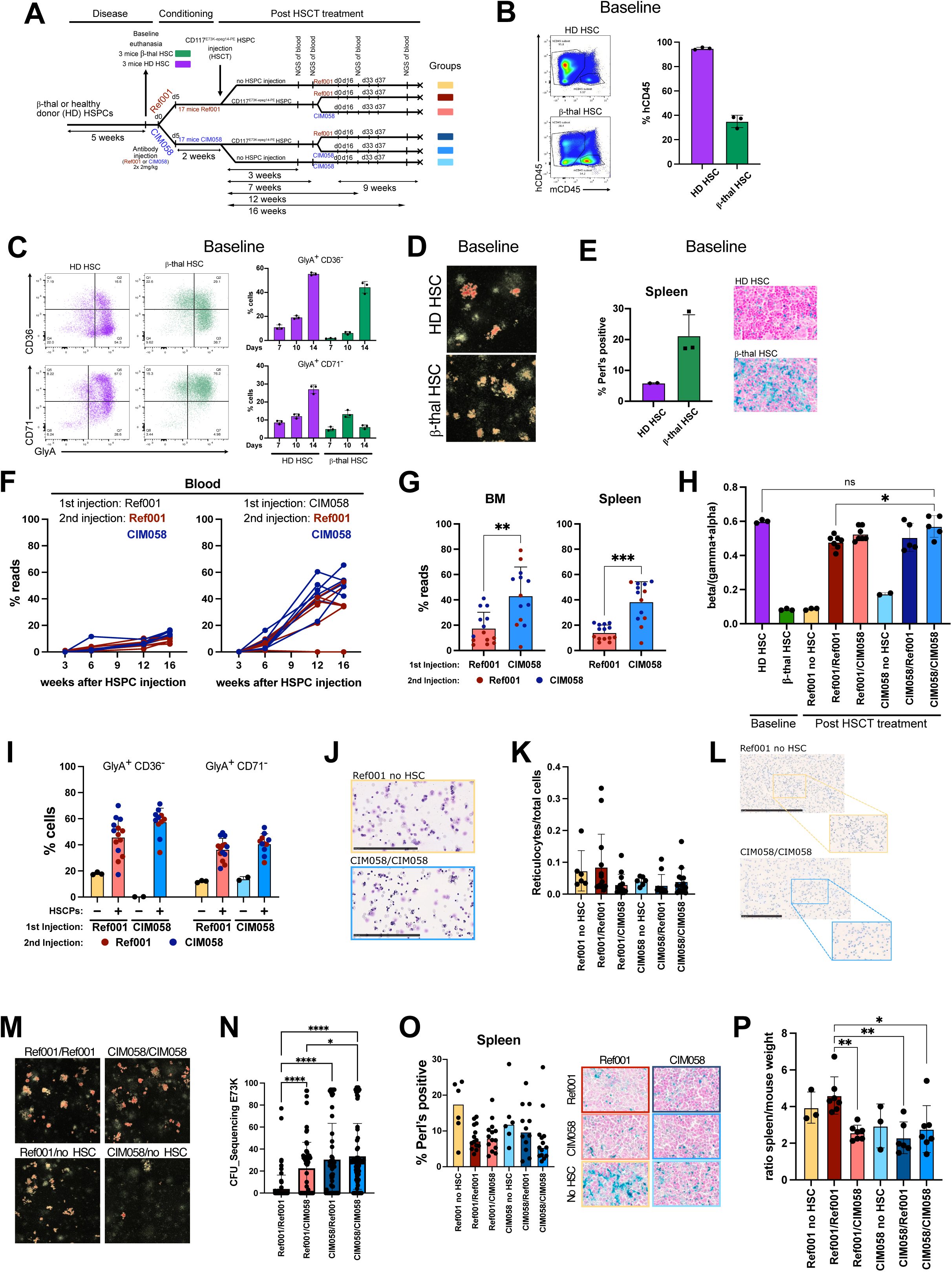
Shielded HSPCs paired with CIM058 ameliorated disease phenotype in a humanized β-thalassemia mouse model (A) Schematic representation of the mouse in vivo experiment. NBSGW mice were engrafted with healthy donor (HD) or beta thalassemic HSPCs. Six week later some mice were euthanised in order to evaluate the baseline engraftment (HD: purple group; β-thal: green group). The other mice received 2 Ref001 or CIM058 injections. Afterwards, the mice received CD117^E73K-epeg14-PE^ HSPC (control mice didn’t received HSPCs) followed by 4 Ref001 or CIM058 injections. (B) Human hematopoietic engraftment at baseline. Representative Flow cytometry plots and quantification of human CD45^+^ in the BM (n = 3 mice per group). (C) Impaired erythroid maturation in β-thal mice. Representative flow cytometry plots and quantification of in vitro erythroid differentiation of BM cells isolated from the mice. Frequency of the %GlyA^+^/CD36^-^ cells (top) and frequency of %GlyA^+^/CD71^-^ cells (bottom) (n = 3 mice per group). (D) Representative picture of BFU-E from BM cells isolated from the mice and plated for 2 weeks in methylcellulose. (E) Perl’s Prussian blue staining of spleen tissue sections. Quantification of iron deposition using a QuPath pipeline (n = 3 mice per group, left panel) and representative pictures (n = 3 mice per group). (F) Longitudinal NGS analysis of the edited reads in blood. Left, mice initially conditioned with Ref001; right, mice initially conditioned with CIM058. In each panel, red lines indicate the mice that received Ref001 after transplantation and blue lines indicate the mice that received CIM058 after transplantation, in subsequent injections. (n = 7 mice per group). (G) NGS quantification of the edited reads in the BM and spleen of the mice at the end of the experiment. Bars indicate the initial conditioning antibody (red, Ref001; blue, CIM058), while dots indicate the post-transplant Ab treatment (red, Ref001; blue, CIM058). (n = 7 mice per group). (H) HPLC analysis of globin chain composition cells following in vitro erythroid differentiation. Quantification of the β/(γ+α)-globin ratio (n = 2-7 mice per group). (I) Flow cytometry analysis of BM cells at the end of erythroid differentiation. Frequency of the %GlyA^+^/CD36^-^ cells (left) and % GlyA^+^/CD71^-^ cells (right). Bars indicate the initial conditioning antibody (red, Ref001; blue, CIM058), while dots indicate the post-transplant Ab treatment (red, Ref001; blue, CIM058). (n = 2-7 mice per group). (J) Representative May-Grünwald/Giemsa-stained cytospins of BM cell erythroid differentiation cultures from untreated β-thal mice (Ref001/no HSCT) and mice receiving CIM058 conditioning, CD117^E73K-epeg14-PE^ HSPC transplantation and post-transplant CIM058 selection. Scale bar = 250 μm (K) Quantification of reticulocytes in peripheral blood smears using a QuPath pipeline (n = 3-7 mice per group, 1-2 slides per mouse). (L) Representative New Methylene Blue staining of peripheral blood of reticulocyte. Reduced reticulocytosis in CIM058/ HSCT/CIM058 mice relative to untreated β-thal controls (Ref001/no HSCT). Scale bar = 250 μm. (M) Representative BFU-E colonies generated from BM cells cultured in methylcellulose for 2 weeks. (N) Sanger sequencing of individual erythroid colonies picked from the CFU assay. E73K reads are plotted. (O) Splenic iron deposition at study endpoint. Quantification of Perl’s Prussian blue staining using a QuPath pipeline (left) and representative spleen sections (right) (n = 3-7 mice per group, 1-2 slides per mouse). (P) Spleen-to-body weight ratio as a measure of splenomegaly (n = 3–7 mice per group). Statistics: Ordinary two-way analysis of variance (ANOVA) tests (significance level, α = 0.05) or Welch’s t test were used to assess statistical difference between groups. Bars represent mean ± SD. Each dot denotes an individual mouse. Significance: *p < 0.05, **p < 0.01, ***p < 0.001, ****p < 0.0001; ns, not significant.

As expected at baseline, β-thal CD34^+^ cells displayed impaired engraftment in comparison to the HD HSPCs, (Figure 6B). Consistent with ineffective erythropoiesis, erythroid differentiation of β-thal BM cells was characterized by delayed acquisition of the late erythroid marker GlyA (CD235a), prolonged expression of the intermediate erythroid marker CD36 and impaired downregulation of the early erythroid marker CD71, compared with HD controls (Figure 6C). In line with these results, colony forming assays from β-thal mice produced paler erythroid colonies over their HD counterparts, indicative of reduced hemoglobinization (Figure 6D), while Perls’ Prussian blue staining demonstrated increased splenic iron deposition (Figure 6E). Collectively, these baseline findings established a disease model recapitulating key pathological features of β-thalassemia.

Next, we tested in the β-thal mice transplanted with the therapeutic allogeneic CD117^E73K-epeg14-PE^ cells whether CIM058 treatment selectively enriched shielded cells. We longitudinally quantified in blood E73K editing by NGS post HSCT. In CIM058 treatment groups (CIM058/Ref 001and CIM058/CIM058), E73K reads, increased progressively up to 60% over the 16 week study period, whereas edited frequencies remained below 20% in mice receiving shielded cells without previous mAb conditioning (Ref001/Ref001 and Ref001/CIM058). Post-transplant selection without previous mAb conditioning (Ref001/CIM058) only resulted in an upward trend of editing (Figure 6F). A similar but more pronounced enrichment was also detected in BM and spleen at study endpoint (Figure 6G), likely reflecting the delay between the depletion of unedited cells in the BM and the subsequent appearance of selected cells in the blood. Across all tissues, mice conditioned with CIM058 (CIM058/Ref001 and CIM058/CIM058) exhibited substantially higher editing rates than mice initially receiving Ref001 (Ref001/Ref001 and Ref001/CIM058) (Figures 6F and 6G), indicating that CIM058 conditioning effectively facilitated engraftment of the incoming HSPCs. In vivo selection with CIM058 in non-conditioned mice (Ref001/CIM058) also enriched E73K reads relative to the Ref001 only group (Ref001/Ref001), although at a lesser extent, illustrating that late selection worked as well.

To assess therapeutic efficacy, we first quantified the beta/(gamma + alpha) globin chain ratio. As expected, this ratio was markedly reduced in the β-thal mice at baseline (Figure 6H) and remained unchanged in mice receiving Ref001 isotype control (yellow bar) or CIM058 (light blue bar) without HSCT. In contrast, transplantation of CD117^E73K-epeg14-PE^ HSPCs improved globin chain balance, most markedly in mice that received CIM058 before and after CD117^E73K-epeg14-PE^ HSPC transplantation (CIM058/CIM058), restoring the ratio to levels comparable to mice engrafted with HD HSPCs and thus demonstrating therapeutic benefit (Figure 6H).

Likewise, impaired erythroid differentiation (Figure 6C) was ameliorated by transplantation of CD117^E73K-epeg14-PE^ HSPCs and further boosted by CIM058 treatment (Figure 6I). This was also reflected in an improved representation of maturing erythroid cells in cytospin samples from CIM058-treated mice at the end of erythroid differentiation culture (Figure 6J). In addition, CIM058 treatment, before and after CD117^E73K-epeg14-PE^ HSPC transplantation, reduced reticulocytes (Figures 6K and 6L) and further enhanced hemoglobinization of erythroid colonies (Figure 6M). Colonies from β-thal mice (Ref001 no HSC), or from mice that received CD117^E73K-epeg14-PE^ HSPCs and Ref001 as treatment (Ref001/Ref001), appeared paler than colonies from mice that received CIM058 before and after HSPC injection (CIM058/CIM058), confirming improved hemoglobinisation (Figure 6M). Mice that received only CIM058, but no edited HSPCs (CIM058/no HSC), formed very few colonies, likely reflecting efficient depletion of β-thal HSPCs. Sanger sequencing of individual erythroid colonies confirmed selective enrichment of edited cells following CIM058 treatment (Figure 6N). Mice receiving CIM058 after transplantation (Ref001/CIM058) displayed higher frequencies of E73K-edited colonies than Ref001/Ref001 controls, while mice conditioned with CIM058 before HSCT (CIM058/Ref001) or/and receiving CIM058 post-transplant (CIM058/CIM058) contained a population of E73K homozygously edited colonies which was absent or very rare in mice that were first treated with isotype control (Ref001/Ref001 and Ref001/CIM058, respectively).

Functional benefit was further supported by improvements in additional disease parameters. Transplantation of CD117^E73K-epeg14-PE^ HSPCs alone (Ref001/Ref001) reduced splenic iron overload compared to untreated β-thal mice (Ref001/no HSC), whereas CIM058 administration before and after HSCT (CIM058/CIM058) produced the greatest reduction on iron burden (Figure 6O). Finally, splenomegaly represents another hallmark of β-thalassemia. We set spleen size of mice engrafted with β-thal CD34^+^ cells as the comparator (Ref001 no HSC). Transplantation of CD117^E73K-epeg14-PE^ HSPCs alone (Ref001/Ref001) failed to reduce the spleen-to-body weight ratio (Figure 6P), whereas CIM058 treatment applied before HSCT with CD117^E73K-epeg14-PE^ HSPCs (CIM058/Ref001), after HSCT (Ref001/CIM058) or at both time points (CIM058/CIM058) significantly reduced spleen size.

Together, these findings demonstrate that CIM058-mediated conditioning and post-transplant selection efficiently enriched CD117-shielded HSPCs (carrying a WT globin allele) in vivo, and substantially improved multiple hematologic hallmarks of β-thalassemia.

## DISCUSSION

In this study, we establish CD117 epitope shielding as a strategy to uncouple targeted depletion of endogenous HSPCs from donor HSPC engraftment. We identified a CD117 variant that preserves physiological receptor function while conferring resistance to a highly potent CD117-depleting mAb, enabling efficient Ab-mediated conditioning without compromising transplanted HSPCs and also permitting sustained post-transplant mAb administration for in vivo selection of shielded cells. This approach has potential applications in both allogeneic HSCT or HSPC gene therapy.

CD117 has long been recognized as a suitable surface receptor for chemotherapy-free conditioning due to its expression by HSPCs^30^. While preclinical studies in mice and NHP supported the feasibility of CD117-targeted conditioning^26,30–32,49^, clinical translation faced setbacks in balancing efficacy and safety. The Fc active, CD117-targeting antibody-drug conjugate (ADC), LOP628, caused hypersensitivity reactions (HSR)^43^ whereas Fc inactive CD117-targeting conventional mAbs (briquilimab and barzolvolimab) were clinically safe^34,50–53^. However, briquilimab monotherapy lacked potency for HSCT conditioning^52^. Conversely, a CD117-ADC, i.e. high potency modality, was associated with a patient death^26^, illustrating the potential risks of ADC-based conditioning. Although alternative payloads and/or conjugation methods might improve safety^26,54^, the conundrum to achieve safe conditioning using an ADC may remain since most ADCs retain some degree of payload-related toxicity^54^. Together, these clinical results highlight the ongoing challenge of achieving effective HSPC depletion without compromising safety.

To overcome the limited potency of SCF-blockade alone for sufficient long-term HSPC depletion and conditioning, while avoiding the necessity for ADCs, we engineered the CD117^E73K^ variant into HSPCs to shield them from depletion by a potent CD117-targeting mAb (CIM058). The shielded HSPCs tolerated repetitive post-HSCT CIM058 administration, enabling sustained depletion of remaining endogenous or non-edited CD117^WT^ cells and progressive in vivo enrichment of engineered cells. This strategy addresses an unavoidable challenge of all potent CD117-targeting cell depleters: the requirement to reduce exposure to the depleter prior to HSPC infusion to prevent donor cell depletion. For CD117-targeting ADCs, previous strategies included engineering a short ADC half-life^26,49^. Although this spares the incoming HSPCs, it requires maximal efficacy to be achieved with a single ADC dose, inherently increasing toxicity risk. In contrast, the CD117 cell shielding approach presented here permits dose-fractionation through repeated administration of the applied depleter and therefore reducing maximal drug exposure which is associated with toxicity. Combining shielded cells with ADCs could therefore reduce the toxicity of peak doses as well as on-target toxicity related to the deep HSPC depletion which typically requires supportive care in NHP^26,55^. Therefore, CD117 shielding in the context of ADCs improves safety. In the β-thalassemia model, the strongest therapeutic benefits were observed in several assays in the group with the most CIM058 applications, before and after HSCT, highlighting the advantage of prolonged post-transplant selection to increase potency. This is particularly relevant because in the absence of cell shielding, briquilimab displayed a maximal dose above which chimerism actually decreased^33,34^. CD117-shielded HSPCs therefore increase the tolerated dose and thus expand the therapeutic window of SCF blocking antibodies and remove the need for a highly potent but short half-life depleter. In support of this conclusion, preliminary results of an independent, ongoing NHP study demonstrated that a toxin-free CD117 blocking mAb was sufficient for conditioning for an autologous HSCT^56^. The transplanted HSPCs were multiplex based edited ex vivo for CD117 shielding and to induce fetal hemoglobin (HbF). CD117-shielded and HbF expressing cells were co-selected in vivo through continuous CD117-blocking mAb administration. Two aspects were remarkable: i) antibody-based conditioning as single agent was sufficient in this immunocompetent large animal model and ii) reaching therapeutic HbF levels was achieved without any supportive care. Together with our data, these results largely support our hypothesis that CD117 HSPC shielding combined with a toxin-free CD117 blocking antibody may provide a clinically feasible alternative approach to conventional genotoxic HSCT conditioning regimens. However, the key remaining open question is the durability of the achieved donor chimerism after discontinuation of antibody treatment. Durable engraftment would indicate that HSPCs were truly eliminated and stable niche replacement occured. Although, we did observe stable enrichment of shielded cells in immunodeficient mice engrafted with human CD34^+^ cells, anti-mouse CD117 antibody conditioning has historically been less effective in immunocompetent mice^30^. Likewise, observations in humans showing recovery of endogenous HSPCs following clearance of briquilimab suggest that SCF blockade alone may transiently displace or suppress rather than permanently eliminate, host HSPCs. Whether durable chimerism can be sustained following cessation of conventional antibody treatment, therefore remains a critical question that will require evaluation in NHP studies and future clinical trials.

Although CD117 HSPC shielding combined with paired mAb infusions may represent the mildest possible conditioning by reducing or eliminating critical risks associated with HSCT, this benefit needs to be carefully weighed against the safety of the engineered cells themselves. In the direct comparison of engineering CD117^E73K^ using CRISPR/Cas9 nuclease-mediated HDR or PE, the latter offered multiple advantages, including highly precise on-target editing without detectable unintended indels or large chromosomal rearrangements and superior engraftment. This was likely not only due to the absence of a DNA HDR template which typically causes cellular toxicity but also related to the reduced genotoxic stress. Although longterm cellular health cannot be quantified e.g. by viability assays, the superior in vivo results must reflect a generally improved fitness of prime-edited HSPCs. Our initial safety assessment using CAST-Seq - although favorable - does not reflect a full safety assessment required for clinical translation. Importantly, PE continues to evolve, and newer improvements such as enhanced RT or epegRNA designs may further increase editing efficiency^57^. A comprehensive preclinical safety evaluation will therefore be most informative once editing conditions have been fully optimized. In fact, in an industrial setting PE for CD117 shielding reached 98% efficiency^58^ while maintaining a highly favorable safety profile even at these high on-target editing rates^59^. Ultimately, the risk/benefit assessment of using engineered cells to enable toxin-free conditioning should be evaluated against the known genotoxicity of conditioning regimens used in current clinical practice.

Lastly, CD117 shielding could have distinct but complimentary applications in autologous HSPC gene therapy and allogeneic HSCT. Autologous HSPC gene therapy is preferred for genetic treatments because it is donor-independent and it eliminates the risks of graft rejection and graft versus host disease (GVHD) while enabling durable hematopoietic reconstitution from the patient’s own cells. Clinical successes in β-thalassemia and SCD have firmly established the feasibility of this approach^19,20,29^. In this setting, CD117 shielding provides the potential advantage to eliminate the need for busulfan-based conditioning. Multiplex genome editing to introduce both the therapeutic edit and the shielding mutation were demonstrated in preclinical studies^56,58^. However, besides the immunological advantages, an autologous co-selection approach has several disadvantages, including increased manufacturing complexity and regulatory burden. Moreover, patient HSPCs may be fragile and multiple apheresis cycles may be necessary. Most importantly, co-selection requires a dedicated therapeutic editor for each mutation which severely constrains its use and is associated with very high development costs. In contrast, the most important advantage of CD117 shielding for the allogeneic setting, in addition to the healthy starting material, is the eliminated need for disease-specific genome correction, since shielded HSPCs carry a functional wild-type allele for any disease indication. The allogeneic approach therefore only requires the development and clinical validation of a single edit, the shielding mutation, thus improving product consistency and affordability. This is particularly attractive for genetically heterogeneous disorders, including primary immunodeficiencies, bone marrow failure syndromes, metabolic disorders, and many other monogenic diseases, where each disease - and often each mutation - requires its own corrective editing strategy. CD117 shielding could be integrated to existing clinical protocols for haploidential HSCT, e.g. for SCD, to reduce toxicity of the currently used conditioning regimen^60^. Although, in allogeneic HSCT GVHD and graft rejection represent safety risks, they can be mitigated by the transplantation of purified, shielded CD34^+^ cells with minimal T cells and by the non-toxic conditioning which reduces tissue damage and the inflammatory signals that contribute to GVHD pathogenesis^61^.

Finally, β-thalassemia illustrates the dynamic nature of therapeutic options being developed for monogenic blood diseases. Shielded allogeneic HSPCs could be attractive for β-thalassemia with >300 known causal mutations^62^, although in this case, HbF reactivation with multiplexed CD117 shielding could provide a mutation-agnostic autologous alternative. However, both options could ultimately co-exist because historically, for non-scientific reasons, even some approved drugs are not available in all regions. Taken together, CD117 shielding represents a platform technology that addresses distinct unmet needs. In autologous gene therapy, it complements therapeutic genome editing by enabling selective in vivo expansion of corrected HSPCs, potentially lowering the editing threshold required for durable clinical benefit. In allogeneic HSCT, it simplifies manufacturing and replaces genotoxic conditioning with a targeted, chemotherapy-free alternative, making this approach safer and broadly applicable across HSCT-treatable diseases. In the future, CD117 shielding may also offer the possibility to enrich cells after in vivo engineering where gene editing efficiencies are currently very low. Rather than favoring one modality, we propose that CD117 shielding could be applied to autologous or allogeneic stem cell therapies or even in vivo gene engineering, depending on availability and needs.

### Limitations of the study

Although our data provide evidence that toxin-free HSCT with CD117-shielded HSPCs carrying a wild-type β-globin gene ameliorates the β-thalassemic phenotype, several limitations of this study should be noted. First, our results represents a scientific proof-of-concept but additional experiments are required for clinical translation. For instance, since this is a highly dynamic field, alternative variants besides CD117^E73K^ may be considered, the PE architecture used to install the variant can likely be further optimized and ultimately a more comprehensive safety assessment should be performed according to applicable regulatory guidelines^63,64^, which are rapidly evolving as well. Second, although immunodeficient mice engrafted with human CD34^+^ cells represent a gold standard in the field, these models also have substantial shortcomings. As an example, the NBSGW mouse strain used for the primary transplants strongly skews differentiation of engrafted human CD34^+^ cells toward the B-cell lineage, which limits the assessment of the potential of engineered cells to differentiate into all hematopoietic lineages. Nevertheless, secondary transplantation experiments, thoroughly support the conclusion that the engineered CD34^+^ cells retain LT-HSC potential. Another limitation particularly relevant for the β-thalassemia experiments is that the NBSGW strain permits engraftment of human CD34^+^ cells without irradiation or other conditioning. Therefore, CD117-shielded HSPCs engrafted even in mice that had not received CIM058, leading to a partial rescue of the β-thalassemic phenotype independent of antibody treatment. This should be born in mind for the interpretation of the results of the β-thalassemia model, although the results still clearly support a beneficial therapeutic effect of CIM058 treatment and enrichment of shielded cells.

## Supporting information

Table antibody

Table guides

Table primers

## RESOURCE AVAILABILITY

### Lead contact

Requests for further information and resources should be directed to and will be fulfilled by the lead contact, Lukas T. Jeker, at

### Materials availability

All noncommercial materials developed in this manuscript are freely available unless a material transfer agreement prevents us from distributing material.

### Data and code availability

CAST-Seq data are available on GEO: GSE336353

## ACKNOWLEDGEMENTS

We thank the members of our laboratories for fruitful discussions and suggestions throughout this work, Alessandro dell’ Aglio, Giuseppina Capoferri, Corinne Engdahl, Hanna Studer, Théa Duchâtel, Anna Haydn, Marcel Heugel, Melanie Hug, Sophie Lackner, Andreja Rasmussen, Olivia Rudin and Mathilde Testut for technical assistance, Robert Ivanek and Florian Geier for bioinformatic support and Julian Grünewald for initial advice with prime editing. We acknowledge the contributions of the following research core facilities at the University of Basel and the Department of Biomedicine: the teams of the animal facility for expert animal husbandry, the bioinformatics core facility, the histology core facility and the flow cytometry core facility for excellent support. Calculations were performed at sciCORE (http://scicore.unibas.ch/) scientific computing center at University of Basel. Epitope mapping was performed at Integral Molecular.

This project has received funding from the European Research Council (ERC) under the European Union’s Horizon 2020 research and innovation programme (grant agreement No. 818806 to L.T.J.) and institutional funds by the Department of Biomedicine (L.T.J.). D.R.L. acknowledges US NIH grants R35GM118062 and RM1HG009490, and the Howard Hughes Medical Institute. We also acknowledge funding from the German Federal Ministry of Education and Research (BMBF) EkoEstMed–FKZ 01ZZ2015 (G.A.).

## AUTHOR CONTRIBUTIONS

Conceptualization: R.M., R.L., S.U. and L.T.J.

Methodology: R.M., R.L., K.P., J.Z., A.S., T.B., M.T., M.R., G.A., A.W. and L.C.W.

Investigation: R.M., R.L., K.P., J.Z., A.S., A.C., T.B., E.B., M.T., J.W., M.R., G.A., C.L., A.H., A.W., L.C.W., E-M.G., E.T.B., J.B., L.G.P., F.L., V.D.S. and C.D.

Formal analysis: R.M., R.L., K.P., J.Z., A.S., A.C., E.B., M.R., G.A., J.B., L.G.P. and F.L.

Data curation: R.M., R.L., E.B., M.R. and G.A.

Software and computational tools: R.L., E.B., M.R. and G.A.

Visualisation: R.M., R.L., K.P., J.Z., A.C., T.B., M.R., A.W., L.C.W., E-M.G., E.T.B., J.B., L.G.P., F.L. and V.D.S.

Writing original draft: R.M. and L.T.J. Writing review and editing: all authors

Supervision: R.M., A.S., D.C., L.G.P., S.Y., D.R.L., A.L., T.I.C., T.C., E.Y., S.U. and L.T.J.

Funding acquisition: L.T.J.

Project administration: S.U., L.T.J.

## DECLARATION OF INTERESTS

Funding: Research was supported by the European Research Council (L.T.J.). Sponsored research agreement with Cimeio Therapeutics AG (Cimeio) (L.T.J. and T.I.C.). Decision to publish was the sole responsibility of L.T.J. L.T.J.’s employer, the Basel University Hospital, receives financial compensation for L.T.J.’s consulting.

Employment: Ridgeline Discovery GmbH: A. S., A. W., L.C.W.

Employment: Cimeio Therapeutics: A. C., E-M. G., E. TB., J. B., L. G. P., F. L., V. DS., C. D., S. Y., S. U.

Cimeio co-founder: R.M., R.L., L.T.J.

Personal financial interests: L.T.J.: co-founder, board member of Cimeio AG.

Holding Cimeio equity: University of Basel, R.M., R.L., A. S., A. C., T. B., A. W., L.C.W., E-M. G., E. TB., J. B., L. G. P., F. L., V. DS., C. D., S. Y., S. U., L.T.J.

Inventors on a patent application related to the findings reported here: R.M., R.L., A. S., A. C., A. W., L.C.W., E. TB., L. G. P., F. L., S. U., L.T.J.

Hold a patent on CAST-Seq (US11319580B2): T.C. and G.A.

D.R.L. is a co-founder of Beam Therapeutics, Prime Medicine, Pairwise Plants, Editas Medicine, and nChroma Bio, companies that use or deliver genome editing or epigenome-modifying agents.

## EXPERIMENTAL MODEL AND STUDY PARTICIPANT DETAILS

### Human HSPCs isolation

Leukopaks were purchased from CytoCare and hCD34^+^ HSPCs were isolated using a proprietary process using CliniMACS Prodigy (Miltenyi). Isolated hCD34^+^ HSPCs were frozen in Cryostor CS5 and stored in liquid N_2_. Cells were thawed and grown in HSPC medium for two days until electroporation (StemSpan SFEM II (StemCell, 09655) supplemented with 100 ng/ml human stem cell factor (hSCF) (Miltenyi, 130-096-695), 100 ng/ml human FMS-like tyrosine kinase ligand (hFlt3)-ligand (Miltenyi, 130-096-479), 100 ng/ml human thrombopoietin (hTPO) (Miltenyi, 130-095-752) and 60 ng/ml hIL-3 (Miltenyi, 130-095-069).

### Mouse husbandry

All animal work was done in accordance with the federal and cantonal laws of Switzerland. Protocols were approved by the Animal Research Commission of the Canton of Basel-Stadt, Switzerland. All mice were housed in a specific pathogen-free condition in accordance with institutional guidelines and ethical regulations. NBSGW (stock 026622) female mice were purchased from Jackson Laboratories. For secondary transplant, NSG–SGM3 female mice (stock 013062) were purchased from Jackson Laboratories.

## METHOD DETAILS

### Epitope mapping and variant selection

An alanine scan was performed via shotgun mutagenesis epitope mapping (Integral Molecular, Philadelphia/PA, USA) as previously described^41^. Briefly, a mutation library of kinase-deficient human CD117 was created by high-throughput, site-directed mutagenesis. Residues of the extracellular domain of CD117 were individually mutated to alanine, with alanine codons mutated to serine, achieving >90% library coverage. The mutant library was arrayed in microplates and transiently transfected into HEK293T cells. Following transfection, cells were incubated with SR-1 antibody (Fab format) under high stringency conditions at concentrations pre-determined using an independent immunofluorescence titration curve on wild-type CD117 and detected using an Alexa Fluor 488-conjugated secondary antibody (Jackson ImmunoResearch #109-546-006). Mutated residues were identified as being critical to the antibody epitope if they did not support the reactivity of the test antibody but did support the reactivity of the reference antibody (antibody YB5.B8 (Invitrogen, Cat. no. 14-1179-82)). Binding of each antibody to each mutant clone was determined in duplicates. For each data point, background fluorescence was subtracted from the raw data, which were then normalized to antibody reactivity with wild type CD117. To identify preliminary primary critical clones, a threshold of >70% wild-type binding to control antibody and <20% wild-type binding to test antibody was applied.

In a second step, each critical residue was subject to comprehensive mutagenesis to selected biophysically appropriate non-alanine amino acids, based on sequence, structure-related and general antibody binding prerequisites of the substituted amino acid. Structural models of CD117 in complex with SCF, based on the available crystal structure (PDB ID: 2E9W), were used to support the in silico analysis of epitope residues and shielding variants. Binding analysis and evaluation of SR-1 antibody binding (hIgG1 format) to each mutant clone were performed as described above with the exception that anti-human IgG secondary antibody was used (Jackson ImmunoResearch, #109-545-003).

### BLI measurement of binding

All BLI measurements were performed either on an Octet Red96e (ForteBio) or on an Octet R8 (Sartorius) at 25 °C with shaking at 1,000 rpm using 1x kinetic buffer (Sartorius, PN: 18-1105). The ECD of CD117 WT and variants were produced and purified by Icosagen.

First, the CD117 variants were screened for their ability to bind the SR-1, CIM001 or CIM058 using three different concentrations of analyte. The antibody was captured by Anti-Human Fc capture biosensor (AHC) (Sartorius, PN: 18-5060) for 300 s at 0.5 to 1 ug/ml. Human CD117 WT and variants, containing only domains 1, 2 and 3 (CD117 D1-2-3) were used as analytes. The analytes were titrated at 500 nM, 50 nM and 5 nM. Association and dissociation of the analyte to antibody was monitored for 300 s and 900 s, respectively. Reference subtraction was performed against buffer only wells. AHC tips were regenerated using 10 mM Gly-HCl pH 1.7. Data were analyzed using the Octet Data Analysis software HT 12.0. Data were fitted (when possible) to a 1:1 binding model. Kinetic rates ka and kd were globally fitted. In a qualitative binding analysis, binding level of the CD117 variants were compared to wild type binding levels and were calculated as percentage. For this qualitative analysis, the binding level of the top analyte concentration at end of the association step was used.

### Thermal stability analysis

Different biophysical properties of the CD117 variants were analyzed. Aggregation and yield of the CD117 variants were determined by size exclusion chromatography (SEC) and dividing the amount of protein after purification by the production volume, respectively. Melting temperature was measured by differential scanning fluorimetry using Sypro Orange (Sigma, PN: S5692-50ul) and an RT-PCR machine (C1000 Thermal Cycler, Biorad). The samples were measured at 0.25-1.0 mg/ml in triplicates in PBS buffer in duplicates and Sypro Orange was added at a final concentration of 5x. The temperature was sequentially increased (0.5 °C each 10 s) from 25 to 95 °C. The fluorescent increase was monitored as a function of temperature and the melting temperature ™ was determined as the inflexion point of the sigmoidal curve and compared to CD117 wild-type. The temperature of protein unfolding transition (Tm) was calculated using the first derivative method.

### Genomic DNA extraction, PCR and Sanger sequencing

All primers are listed in the Supplementary Table Primers.

Genomic DNA of cells was extracted using QuickExtract (Lucigen, QE09050). Cell pellets were resuspended in 30-50 µl QuickExtract, incubated at 65 °C for 6 min, vortexed for 1 min and subsequently re-incubated at 98 °C for 10 min. PCR was performed using GoTaq G2 Green Master Mix (Promega, M782B). The gDNA of samples analysed by NGS was extracted using QuickExtract (Lucigen, QE09050) or the Quick-DNA 96 Plus kit (Zymo, D4070) and the genomic DNA concentration was measured with a Qubit device (Thermo Fisher). For sequencing, different PCR primers were used depending on sequencing technology (Supplementary Table). Sequencing of PCR amplicons was done at Microsynth and sequencing chromatograms were analysed using the MultiEditR package to retrieve editing efficiencies^65,66^.

### Next-generation amplicon sequencing

All primers are listed in the Supplementary Table Primers.

For NGS, targeted amplicon libraries were generated using a three-step PCR protocol as described earlier^37^. In brief, nested PCRs were done on genomic DNA samples using KAPA HiFi HotStart polymerase (Roche). Applied primer sequences are listed in Primer Table. Libraries were paired-end sequenced on an Illumina Miniseq instrument using the Illumina Miniseq Mid output kit (300 cycles) with 50% PhiX spike-in (Illumina). After demultiplexing, each sample was assessed for quality using FastQC and processed using the CRISPResso2 tool (v2.3.1)^67^. The quantification window was expanded to 15 bp and centered 10 bp upstream of the canonical cleavage site (--quantification_window_size 15, --quantification_window_center -10) to capture the full editing window. Prior to alignment, reads were trimmed for Nextera adapters using Trimmomatic (pattern: ILLUMINACLI P:NexteraPE-PE:2:30:10 LEADING:3 TRAILING:3 SLIDINGWINDOW:4:15 MINLEN:36), and paired-end merging was performed with a maximum overlap of 200 bp. For visualization purposes, a plot window of 30 bp was utilized, and only alleles with a frequency ≥0.1% were included in the reported allele frequency plots to focus on major editing outcomes. All other parameters were maintained at default settings.

### Cell line culture conditions

TF-1 was purchased from DSMZ (Cat#ACC334) and maintained in RPMI-1640 media (Cat#A1049101, Gibco) supplemented with 10% heat-inactivated FBS (Cat#P30-1909, PAN), and 2 ng/ml hGM-CSF (Cat#PHC2015, Gibco).

HSPCs culture media: StemSpan AOF (Cat#100-0130, STEMCELL Technologies) supplemented with 100 ng/ml Flt3 ligand (Cat#130-096-479, Miltenyi Biotec), 100 ng/ml TPO (Cat#130-095-752, Miltenyi Biotec), 100 ng/ml SCF (Cat# #130-096-695, Miltenyi Biotec) and 60 ng/ml IL-3 (Cat#130-095-071, Miltenyi Biotec).

High Cytokine media: StemPro™-34 SFM (1x) including nutrient supplements (Cat#10639011, Gibco Life Technologies) supplemented with 20 ng/ml Flt3 ligand (Cat#130-096-479, Miltenyi Biotec), 50 ng/ml TPO (Cat#130-095-752, Miltenyi Biotec), 100 ng/ml SCF (Cat# #130-096-695, Miltenyi Biotec), 10 ng/ml IL-3 (Cat#130-095-071, Miltenyi Biotec), 50 ng/ml IL-6 (Cat#130-093-934, Miltenyi Biotec), 20 ng/ml GM-CSF (Cat#PHC2015, ThermoFisher), 3 ng/ml EPO (Cat#78007, STEMCELL Technologies), 10 ng/ml IL-2 (Cat#130-097-743, Miltenyi Biotec), 20 ng/ml IL-7 (Cat#130-095-363, Miltenyi Biotec), 50 ng/ml IL-11 (Cat#130-103-439, Miltenyi Biotec), 500 ng/ml LDL (Cat#2698, STEMCELL Technologies), 1X penicillin-streptomycin (Cat# 15140122, Gibco Life Technologies) and 1X L-glutamine (Cat# 25030081, Gibco Life Technologies).

Low Cytokine media: StemPro™-34 SFM (1x) including nutrient supplements (Cat#10639011, Gibco Life Technologies) supplemented with 20 ng/ml Flt3 ligand (Cat#130-096-479, Miltenyi Biotec), 50 ng/ml TPO (Cat#130-095-752, Miltenyi Biotec), 100 ng/ml SCF (Cat# #130-096-695, Miltenyi Biotec), 500 ng/ml LDL (Cat#2698, STEMCELL Technologies), 1X penicillin-streptomycin (Cat# 15140122, Gibco Life Technologies) and 1X L-glutamine (Cat# 25030081, Gibco Life Technologies).

### Engineering of TF-1 cells expressing CD117 variants

All the sgRNAs and HDRT are listed in the Supplementary Table Guides.

TF-1 cells (0.2 × 10^6^) were washed twice with PBS and resuspended in 10 µl R buffer (Neon Transfection System). Ribonucleoprotein (RNP) complexes were pre-assembled by mixing sgRNA (Synthego (Modified EZ Scaffold), final concentration 75 pmol) and SpyFi Cas9 nuclease (Aldevron, final concentration 31 pmol) and incubating for 20 min at room temperature in the dark. Single-stranded DNA (ssDNA) HDR templates (IDT (Alt-R HDR modification), final concentration 100 µM) were added to the RNP complexes. The RNP–ssDNA mixture was added to the cells and electroporated using the Neon Transfection System (1200 V, 40 ms, 1 pulse) per the manufacturer’s protocol. Edited cells were flow-sorted based on surface staining for SR-1 and 104D2 to yield SR-1⁻ 104D2⁺ (knock-in) population.

### Binding of SR-1 antibody to TF-1 cells

CD117-expressing cell lines (TF-1 WT and variant forms) were seeded at a density of 1×10^6^ cells/ml in FACS Buffer (PBS + 2% FCS +1mM EDTA). Prior to staining, cells were blocked with Human TruStain FcX™ (BioLegend, Cat# 422302) for 15 min at 4 °C. Serial dilutions of SR-1 antibody were prepared to yield final concentrations ranging from 5000 ng/ml to 0.15 ng/ml, and cells were incubated with the indicated concentrations for 30 min at 4 °C. Cells were then stained with a secondary antibody Goat anti-human IgG (Cat# A11013, Thermo Fisher) for 30 min at 4 °C. 7-AAD viability dye (Cat# 420404, Biolegend) was added for the final 10 min at room temperature (RT) in the dark. Flow cytometric acquisition was performed using a NovoCyte Quanteon (Agilent), and data were acquired and analyzed with NovoExpress software (Agilent).

### SCF binding to TF-1 cells

CD117 expressing cell lines (TF-1 WT, TF-1 E73K clone 1 and 2) were seeded at a density of 1×10^6^ cells/ml in FACS Buffer (PBS + 2% FCS +1mM EDTA). Prior to staining, cells were incubated with Human TruStain FcX™ (Cat# 422302, BioLegend) for 15 min at 4 °C to block Fc receptors. After this incubation, two washing steps in FACS Buffer were performed. Biotinylated human stem cell factor (SCF, Cat#SCF-H82E1, AcroBiosystems, final concentration: 1 µg/ml) was pre-complexed at 1:1 ratio with streptavidin-PE (Cat# SA10041, Invitrogen; final concentration: 1µg/ml) for 45 min at room temperature (RT). Cells were then stained with the pre-complexed mixture for 30 min at 4 °C. 7-AAD viability dye (Cat# 420404, BioLegend) was added for the final 10 min incubation at RT in the dark. Flow cytometric analysis was performed using a NovoCyte Quanteon (Agilent) and data were acquired and analyzed with NovoExpress Software (Agilent).

### SCF dependent proliferation assay of TF-1 cells

SCF dependent proliferation was assessed using TF-1 CD117 expressing cell lines. TF-1 and its variants (E73K and S123F) were seeded at a density of 1.5×10^5^ cells/ml in triplicate in RPMI/FCS medium without GM-CSF and pre-incubated for 1 h at 37 °C and 5% CO₂. Serial dilutions of stem cell factor (SCF; Cat# GMP300-07, PeproTech) were prepared to yield final concentrations ranging from 50 ng/ml to 0.73 ng/ml. Assay plates were incubated with the indicated concentrations for up to 72 h at 37 °C and 5% CO₂. Cell proliferation was quantified using the CellTiter-Glo® 2.0 assay (Cat# 9242, Promega) according to the manufacturer’s protocol.

### SR-1 mediated inhibition of SCF-induced proliferation of TF-1 cells

TF-1 CD117-wild-type and two CD117 variants (E73K and S123F) were seeded at a density of 1.5x 10^5^ cells/ml in triplicate in RPMI/FCS medium without GM-CSF and pre-incubated for 1 h at 37 °C and 5% CO₂. Cells were stimulated with stem cell factor (SCF; final concentration: 100 ng/ml), followed by treatment with SR-1 antibody (Icosagen) at concentrations ranging from 50 µg/ml to 0.098 µg/ml. Plates were incubated for up to 72 h at 37 °C and 5% CO₂. Cell proliferation was quantified using the CellTiter-Glo® 2.0 assay (Promega, Cat# 9242) according to the manufacturer’s instructions.

### c-Kit phosphorylation ELISA

TF-1 cells expressing CD117 WT; E73K variants and CD117 KO were seeded at 2x 10^6^ cells per well in 6-well plates in RPMI/FCS medium without GM-CSF. Cells were treated for 5 min with indicated antibody (final concentration 0.5 µg/ml, SR-1, Icosagen; CIM001, Icosagen) and SCF (final concentration 100 ng/ml, Cat# GMP300-07, PeproTech), or with indicated antibody or SCF alone. For SCF titration the following concentrations were used: 20, 8, 4, 2,

0.5 ng/ml. Cells were harvested by low-speed centrifugation (1200 rpm), washed with ice-cold PBS, and lysed in 450 µl 1× cell lysis buffer supplemented with phosphatase inhibitors (Cat# P5726 and P0044 Sigma-Aldrich) and a protease inhibitor cocktail (Cat# 8340, Sigma-Aldrich). Lysates were flash-frozen in liquid nitrogen and thawed at 37 °C twice. Subsequently, lysates were centrifuged at 14,000 rpm for 10 min at 4 °C. Supernatant was transferred to a new tube. Phospho-c-Kit (Tyr719) levels were quantified using a Sandwich ELISA (Cat# 7298, Cell Signaling Technology) in duplicate according to the manufacturer’s instructions.

Note: Ctrl. = Addition of H2O for SCF or PBS for SR-1

### CRISPR/Cas9 mediated engineering of HSPCs

All the sgRNAs, epegRNAs and HDRT are listed in the Supplementary Table Guides.

For homolgy directed repair screening, gRNAs were freshly prepared as outlined above, but 50 µM crRNA and tracrRNA were used to form the gRNA and complexed with 1 µM Spyfi Cas9 (Aldevron at 61.889 µM) at a molar ratio Cas9:gRNA = 1:2 and incubated for 20 min at RT. As a control, incomplete RNPs lacking the site-specific crRNA were generated. During the RNPs complexing, HSPCs were collected, washed twice with PBS, and resuspended in 100 µl Lonza supplemented P3 electroporation buffer at 1 × 10^6^ cells/90 µl. Cells were then mixed with 5 µl RNP and ssDNA HDRT encoding the variants (5 µl corresponding to 500 pmol), and the whole volume was transferred into the EP nucleocuvette. Cells were electroporated using the 4D-Nucleofector system (Lonza) with program CA-137. Immediately after EP, the cells were transferred to a six-well plate and rested for 20 min at RT. After 20 min, 2 ml of prewarmed HSC medium supplemented with 100 ng/ml SCF, 100 ng/ml TPO, and 100 ng/ml Flt3L was added and the plate was incubated at 37°C.

For HDR at larger scale for mouse production, after a wash in P3 nucleofection solution (Lonza) or MaxCyte® EP buffer, HSPCs were resuspended in the same buffer at a target concentration of 5-10×10^7^/ml with 3-9 µM sgRNA (Synthego) and 1-3 µM Spyfi Cas9 (Aldevron) at a molar ratio Cas9:gRNA = 1:3 pre-complexed for 20 min at RT and 5 min additional with 5-7.5 µM HDRT (IDT), then electroporated with Lonza 4D or MaxCyte GTx^TM^ electroporator devices. Immediately after EP, the cells were transferred into a culture vessel of appropriate size and, 20min later, resuspended at 0.5×10^6^ cells/ml in pre-warmed HSPC medium. After 48h, cells were harvested and frozen in Mr Frosty until injection.

For BE HSPCs cells were thawed and cultured for 20 h in StemSpan medium supplemented with SCF, Flt3L, TPO, IL-3. A total of 2 × 10^5^ cells were electroporated with BE mRNA (100 μg/ml) and sgRNA (5.5 μM) in P3 Primary Cell Solution on a Lonza 4D-Nucleofector System using program CA-137. Following electroporation, cells were cultured for three days in cytokine-supplemented medium (SCF, Flt3L, TPO) before harvesting for gDNA extraction.

For prime editing in small scale, epegRNA, nicking guide and PEmax were mixed together (180 pmol : 120 pmol : 2 µg, respectively) at room temperature (RT). Per reaction, 0.3 × 10^6^ HSPC cells were resuspended in 20 µl Lonza supplemented P3 electroporation buffer. Cells were then mixed with the PE components, and the whole volume was transferred into the EP nucleocuvette. Cells were electroporated using the 4D-Nucleofector system (Lonza) with program DS-130. Immediately after EP, 80 µl prewarmed HSC medium supplemented with 100 ng/ml SCF, 100 ng/ml TPO, and 100 ng/ml Flt3L was added and the cells were incubated for 15 min at room temperature. Afterward, the cells were transferred to a 48-well plate containing 300 µl prewarmed supplemented HSC medium and the plate was incubated at 37°C. For the experiments in which MLH1dn was used, 1 µg of MLH1dn mRNA was mixed with the other PE components. For big scale electroporation, epegRNA, nicking guide and PEmax were mixed together (594 pmol : 396 pmol : 3.2 µg, respectively). Per reaction, 1 × 10^6^ HSPC cells were resuspended in 100 µl Lonza supplemented P3 electroporation buffer, mixed with the PE components and electroporated. Immediately after EP, 500 µl prewarmed supplemented HSC medium was added and after 15 min the cells were transferred to a six-well plate containing 2 ml prewarmed supplemented HSC medium and the plate was incubated at 37°C.

### SCF binding of edited HSPC in the presence of SR-1 and CIM058 antibodies

CD34⁺ hematopoietic stem and progenitor cells (HSPCs) were seeded in Low Cytokine medium (described above) at a density of 1×10^5^ cells/ml and cultured for 4 days at 37 °C and 5% CO₂. In order to analyze SCF binding in the presence of different antibodies, HSPCs were recovered, washed once with PBS and incubated with Human TruStain FcX™ (Cat# 422302, BioLegend) for 15 min at 4 °C. Cells were then treated with either an isotype control antibody (Ref001, Icosagen), SR-1 antibody (Icosagen), or CIM058 antibody (Icosagen) at a final concentration of 50µg/ml for 15 min at 4 °C. Biotinylated human stem cell factor (SCF, Cat#SCF-H82E1, AcroBiosystems, final concentration: 100 µg/ml) was pre-complexed at 1:1 ratio with streptavidin-PE (Cat# SA10041, Invitrogen; final concentration: 100 µg/ml) for 45 min at room temperature (RT). Serial dilutions of pre-complexed mix were prepared to yield final SCF concentrations ranging from 100 ng/ml to 0.14 ng/ml, and cells were stained with the indicated concentrations for 45 min at 4 °C. Fixable viability stain 780 (Cat# 565388, BD Horizon™) was added for the final 10 min at RT in the dark. Flow cytometric acquisition was performed on a NovoCyte Quanteon (Agilent) and data were acquired and analyzed using NovoExpress software (Agilent).

### SR-1 and CIM058 mediated inhibition of SCF-induced proliferation of HSPCs

CD34^+^ HSPCs were seeded at a density of 1.5x 10^5^ cells/ml in triplicate in High Cytokine medium (described above). Cells were pre-incubated for 1 h at 37 °C and 5% CO₂, followed by treatment with either an isotype control antibody (Ref001, Icosagen), SR-1 antibody (Icosagen), or CIM058 (Icosagen) at concentrations ranging from 1000 ng/ml to 0.15 ng/ml. Plates were then incubated for up to 72 h at 37 °C and 5% CO₂. Cell proliferation was quantified using the CellTiter-Glo® 2.0 assay (Promega, Cat# 9242) according to the manufacturer’s instructions.

### Flow cytometry analysis

Flow cytometry was done on BD LSRFortessa instruments with BD FACSDiva software. Data were analysed with FlowJo software. Antibodies used for flow cytometry are listed in Supplementary Table Antibody.

### Methylcellulose CFU assay of HSPCs in the presence of antibodies

To evaluate the inhibition of colony growth of WT CD34^+^ HSPCs by anti-CD117 antibodies, 500 unedited HSPCs were cultured for 4 days in HSPCs culture medium (described above) then seeded into tubes containing 3 ml of MethoCult™ GF H84435 methylcellulose medium (StemCell Technologies) with the indicated concentrations of SR1, CIM001, CIM058 antibodies or isotype control antibody Ref001 (all from Icosagen).

To evaluate the shielding potency *in vitro*, 300 unedited or edited CD34^+^ HSPCs at 48 hours post-EP were plated with 10 µg/ml Ref001 or CIM058 antibodies.

For all tested conditions, cells in methylcellulose were mixed thoroughly, and 1.1 ml was plated per well of a SmartDish (StemCell Technologies) in duplicates. Plates were incubated at 37°C and 5% CO2 with ≥ 95% humidity for 14-16 days. The resulting colonies were counted and scored with STEMvision^TM^ automated counter (StemCell Technologies) as per the manufacturer’s instruction. When indicated, individual colonies were picked, washed with PBS, lysed in 25 µL QuickExtract^TM^ DNA extraction solution (LGC Biosearch Technologies) at 60°C for 10min, 100°C for 10min and finally at 4°C. Genomic DNA from the lysates was used for CD117 amplification and PCR amplification products were sent for Sanger sequencing (Microsynth).

### Methylcellulose CFU assay of bone marrow cells from *in vivo* studies

40 % of the BM cells isolated from each mouse were cryopreserved to perform Colony Forming Unit (CFU) assays for zygosity analysis of the CD117 edit by Sanger sequencing. Each vial of cryopreserved BM cells was thawed in 10 ml of 50% Ex vivo Lonza medium containing 50% of FBS in a drop-wise manner. After centrifugation, cells were counted and resuspended in 0.5 ml of FACS Buffer.

CFU assay with bone marrow unsorted cells: To generate colonies from the bulk of bone marrow (BM) cells, approximately 20.000 and 40.000 cells for each pool of samples were seeded in FACS tubes containing 3 ml of MethoCult™ medium (thawed at 4°C overnight). 1.1ml of cell suspension was dispensed into each well of a Stemcell SmartDishTM. Plates were incubated at 37 °C, in 5% CO2 with ≥ 95% humidity for 14 – 16 days until colony picking.

CFU assay with human CD45^+^CD34^+^ sorted cells from bone marrow samples: Briefly, cells were stained for 15min at RT (mCD45, hCD45, CD34, CD38, CD117 clone SR-1 and CD117 clone 104D2) and washed once with FACS Buffer. After centrifugation, each sample was resuspended in 0.5ml of FACS Buffer containing the viability dye.

The SONY Sorter MA900 was used to sort and seed 300 human CD45^+^ CD34^+^ cells in each tube containing MethoCult™ medium and seeded as above. Plates were incubated at 37 °C, in 5% CO2 with ≥ 95% humidity for 14 – 16 days until colony picking.

### Methylcellulose CFU assay of prime edited HSPCs

CFU assay was started at 72 h after editing. For each condition, 1.1 ml of semi-solid methylcellulose medium (StemCell Technologies) containing 500 cells was plated in a well of a SmartDish (StemCell Technologies) in duplicates. The cells were incubated at 37°C, 5% O2, and 5% CO2 for 14 days. The resulting progenitor colonies were counted and scored with STEMVision analysis (StemCell Technologies) as per the manufacturer’s instruction. Colonies were picked and subjected to sequencing. Shortly, colonies were washed with PBS and then resuspended in 25 µl DNA-QuickExtract solution. PCRs were performed and sent for Sanger sequencing.

### Methylcellulose CFU assay from mice of the beta-thalessemia experiment

To assess the clonogenic capacity of human cells derived from the chimeric bone marrow, 5 × 10^5^ hCD45⁺ cells were diluted in 300 µL AMEM (Corning™) and subsequently seeded in 2 ml methylcellulose medium (MethoCult™ H4434, Stemcell Technologies). Following 14-day incubation, colony-forming unit-granulocyte-monocyte (CFU-GM) and burst-forming unit-erythroid (BFU-E) were counted with STEMVision analysis (StemCell Technologies). Individual CFU-GM and BFU-E colonies were picked and processed for sequencing.

### Differentiation of prime edited HSPCs in vitro

Three days after EP, cells were resuspended in StemPro media (Gibco) containing StemPro Nutrients, low density lipoprotein 50 ng/ml, P/S 1%, glutamine 1%, Flt3 20 ng/ml, TPO 50 ng/ml, IL-650 ng/ml, IL-3 10 ng/ml, IL-2 10 ng/ml, IL-7 20 ng/ml, erythropoietin 3 ng/ml, GM-CSF 20 ng/ml, and SCF 100 ng/ml, and 2,000 cells/well were plated in a round bottom 96-well plate. In some wells, CIM058 antibody at a concentration of 10 ng/ml was added. After 14 d cells were collected, stained for CD33, GlyA/CD235a and acquired on a Fortessa.

### CAST-Seq

Cells were nucleofected with RNP complexes or PE tools to target CD117.

Genomic DNA was extracted 5 d later using the NucleoSpin Tissue kit (Machery and Nagel). CAST-Seq library preparations were performed as described^44^, and data were analyzed using an improved bioinformatics pipeline^68^. A stringent p-value threshold of 0.005 was applied to minimize the risk of false-positive OMT. Applied primer sequences are listed in Supplementary Table Primers.

### Animal experiments

All animal work was done in accordance with the federal and cantonal laws of Switzerland. Protocols were approved by the Animal Research Commission of the Canton of Basel-Stadt, Switzerland. All mice were housed in a specific pathogen-free condition in accordance with institutional guidelines and ethical regulations. NBSGW (stock 026622) female mice were purchased from Jackson Laboratories. HSPCs were edited as described above (HDR and prime editing). Two days after electroporation, cells were collected and frozen in CryoStor CS10 medium. Cells were thawed on the day of injection, washed and resuspended in PBS. Recipient NBSGW female mice (4 weeks old) were injected intravenously into the tail vein with 1 million HSPCs. Chimerism was analysed by flow cytometry or NGS in blood as indicated in the figures. Mice were treated with saline or CIM058 at the dose(s) and intervals indicated in each figure. Primary transplant mice were euthanized, the blood, spleen and BM were isolated and analysed by FACS, NGS and samples were collected for CFU assays or secondary transplants.

For secondary transplant, NSG–SGM3 female mice (stock 013062) were purchased from Jackson Laboratories. Recipient mice were irradiated the day before the BM transplant with 200 cGy. 40% of the BM from the primary transplant was re-injected into the new host. Mice from secondary transplants were euthanized 8 weeks after humanization.

For the β-thalassemia experiment CD34^+^ cells from β-thalassemia patients (n=2 genotype: IVS1-110/IVS1-110) with β-thalassemia were previously collected during mobilization clinical trials conducted at George Papanikolaou Hospital, Thessaloniki, Greece^48^. NBSGW (stock 026622) female mice were purchased from Jackson Laboratories. On the day of transplantation, cryopreserved CD34^+^ cells were thawed, washed, and resuspended in PBS. Recipient NBSGW mice were transplanted via tail vein injection with 1 × 10^6^ HSPCs per mouse. Five weeks after the initial transplantation, the mice received two injections of CIM058. Two weeks later 1 × 10^6^ CD117 prime edited HSPCs were transplanted via tail vain. Mice were treated with saline or CIM058 at the dose(s) and intervals indicated in each figure.

### Tissue collection and processing

After the mice were euthanized, 0.2 ml blood, both hind legs (femur and tibia) and the spleen were collected from each mouse. Cell suspensions were generated, red blood cells were lysed using ACK lysis buffer and then the cell suspensions were filtered. Cells were stained for different antigens and 30 μl CountBright Absolute counting beads (51’000 microspheres per 50 μl; Invitrogen, C36950) were added to each sample and the results were analysed by FACS using a BD LSRFortessa instrument.

For some experiments, the liver and half of the spleen were fixed for 48h in 4% formaldehyde, washed with PBS and then embedded in paraffin by standard protocol.

Blood smears were prepared by placing one drop of blood-heaprin mix onto a glass slide and spread it using a second slide. Slides were then coverslipped and scanned by a Hamamatsu S60 slide scanner at 40X magnification.

For reticulocytes staining 3 drops of blood were thoroughly mixed with two drops of Reticulocyte Stain (Reticulocyte Stain R4132, Sigma Aldrich) and incubated at room temperature for 10 min. One drop was placed onto a glass slide and spread using a second slide. Slides were then mounted and acquired by slide scanner as described above.

Erythroid differentiation of chimeric bone marrow cells from engrafted mice was carried out as previously described in 3 stages^69^. For the initial ex vivo erythroid culture, 5 × 10⁶ hCD45⁺ cells were seeded. During stage I the cells were expanded in the presence of hydrocortisone 1uM, IL-3 5ng/ml, SCF 100ng/ml and EPO 3U/ml, during stage II only in the presence of SCF and EPO and in the final stage III, SCF was omitted. On days 7, 11, 14 and 18, aliquots of cells were collected, and stained with anti-hCD235a, anti-hCD36 and anti-hCD71 and analyzed by flow cytometry.

For cytospin slide preparation, human bone marrow cells from engrafted mice were depleted of mouse cells using mouse CD45 MicroBeads (130-052-301 Miltenyi). A total of 1.5-2 × 10^5^ cells were prepared by cytocentrifugation (Cytospin 4, ThermoScientific) at 300 g for 5 minutes. Cells were stained with May-Grünwald-Giemsa staining, and their morphology was subsequently examined.

### Histological staining

May–Grünwald–Giemsa staining was performed by incubating slides in May–Grünwald stock solution (Carl Roth, ref. T863.1) for 3 min, followed by two washes in water and staining with Giemsa working solution (10% v/v in ddH₂O; Carl Roth, ref. T862.1) for 10 min. Slides were washed twice in water, air-dried overnight at room temperature in an upright position, mounted with coverslips.

For H&E and Perl’s staining, FFPE livers were sectioned at 3 µm and spleens at 5 µm. Sections were incubated at ∼60°C for 10 min, deparaffinized in Ultraclear (3 × 2 min; J.T.Baker, ref. 3905.9010PE), and rehydrated through a graded ethanol series, followed by rinsing in tap water.

For H&E staining, sections were stained with Harris’s hematoxylin (Histolab, ref. 01800-EX) for 1 min, rinsed in running tap water until clear, differentiated in acid alcohol (Leica Biosystems, ref. 3803650E) for 2 min, and blued in Scott’s tap water (Leica Biosystems, ref. 3802901E) for 2 min. Slides were washed in tap water followed by distilled water, counterstained with eosin (CellaVision, ref. 312740) for 4 min, washed again in tap water, dehydrated through graded ethanol, cleared in Ultraclear, and coverslipped.

Perl’s staining was performed using the Prussian Blue Iron Detection Kit (Morphisto, ref. #11097). Sections were deparaffinized and rehydrated as described above, incubated in potassium hexacyanoferrate for 5 min, and subsequently treated with potassium hexacyanoferrate–hydrochloric acid solution (1:1) for 30 min. After washing in water, slides were counterstained with Nuclear Fast Red for 10 min, dehydrated, cleared, and mounted.

For all, slides were scanned by a Hamamatsu S60 slide scanner at 40X magnification.

### HPLC

Individual globin chain levels from transplanted NBSGW bone marrow-engrafted human cells cultured for 18 d in erythroid differentiation media were quantified by reverse-phase HPLC. Globin chains from ex vivo differentiated thalassemic cells were quantified on a Shimadzu LC-2060C 3D Liquid chromatography with a GmbH MultoHigh Bio 300, 250×3 mm column. A 38%-60% gradient mixture of 0.1% trifluoroacetic acid in water/acetonitrile was applied at a rate of 1 ml/min.

### Image analysis

QuPath (v.0.5.1)^70^ was used for image analysis.

For Perl’s staining deposition: Tissue was detected using a pixel thresholder on mean RGB channels, with pixel size= 7.0 µm; gaussian filter and sigma= 1.5.

For iron deposition measurement, an ANN_MLP pixel classifier was trained on a composite image from 8 regions of interest. Iron positive and iron negative regions were manually annotated and the classifier was trained with parameters: pixel size: 0.44µm; channels RGB; filter; gaussian, scales= 1. Percentage of iron staining area was calculated in relation to the total tissue area.

For reticulocytes detection: region of interest for cell detection in a blood smear was manually determined to avoid slide edges and areas without cells. Cell detection was performed using the Watershed algorithm from QuPath with the following parameters: channel for detection= Hematoxylin, pixel size= 0.44µm, threshold=0.05 and no cell expansion. N=20 cells were further manually annotated as Reticulocytes or Others and a Random Trees Object classifier has been trained on all measurements. The number of reticulocytes was expressed as a percentage compared to the total number of detected cells.

## FIGURE LEGENDS

**Figure S1:**
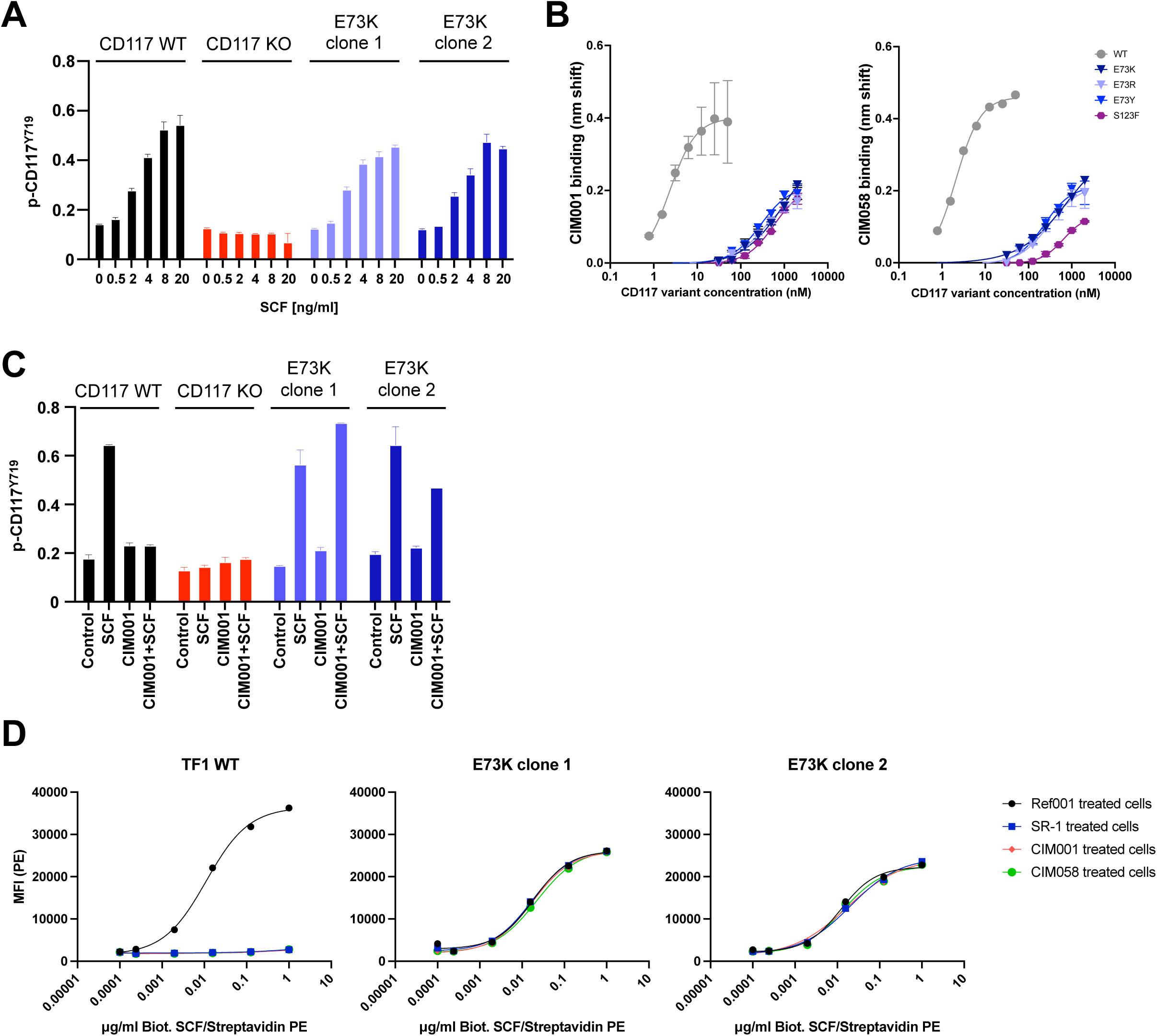
related to Figure 2: CD117^E73K^ retain SCF dependent function despite molecular shielding from SR-1 and derivatives (A) CD117 phosphorylation at position Tyr719 upon increasing SCF concentrations in TF-1 cells expressing wildtype, KO CD117 or the engineered variant E73K. n = 3 biological replicates with 2 technical replicates. (B) Dose titration of selected CD117 variants to test residual CIM001 or CIM058 binding at very high analyte concentrations. (C) CD117 phosphorylation at position Tyr719 upon anti-CD117 antibody treatment in TF-1 cells expressing wildtype, KO CD117 or the engineered variant E73K; SCF: 0.1µg/ml; CIM001: 0.5µg/ml. (D) Binding of the SCF to TF-1 cells expressing engineered CD117 variant E73K in comparison to cells expressing the wildtype receptor in the presence of different CD117 mAbs (SR-1, CIM001 and CIM058). Shown are results from two independent TF-1 E73K clones.

**Figure S2:**
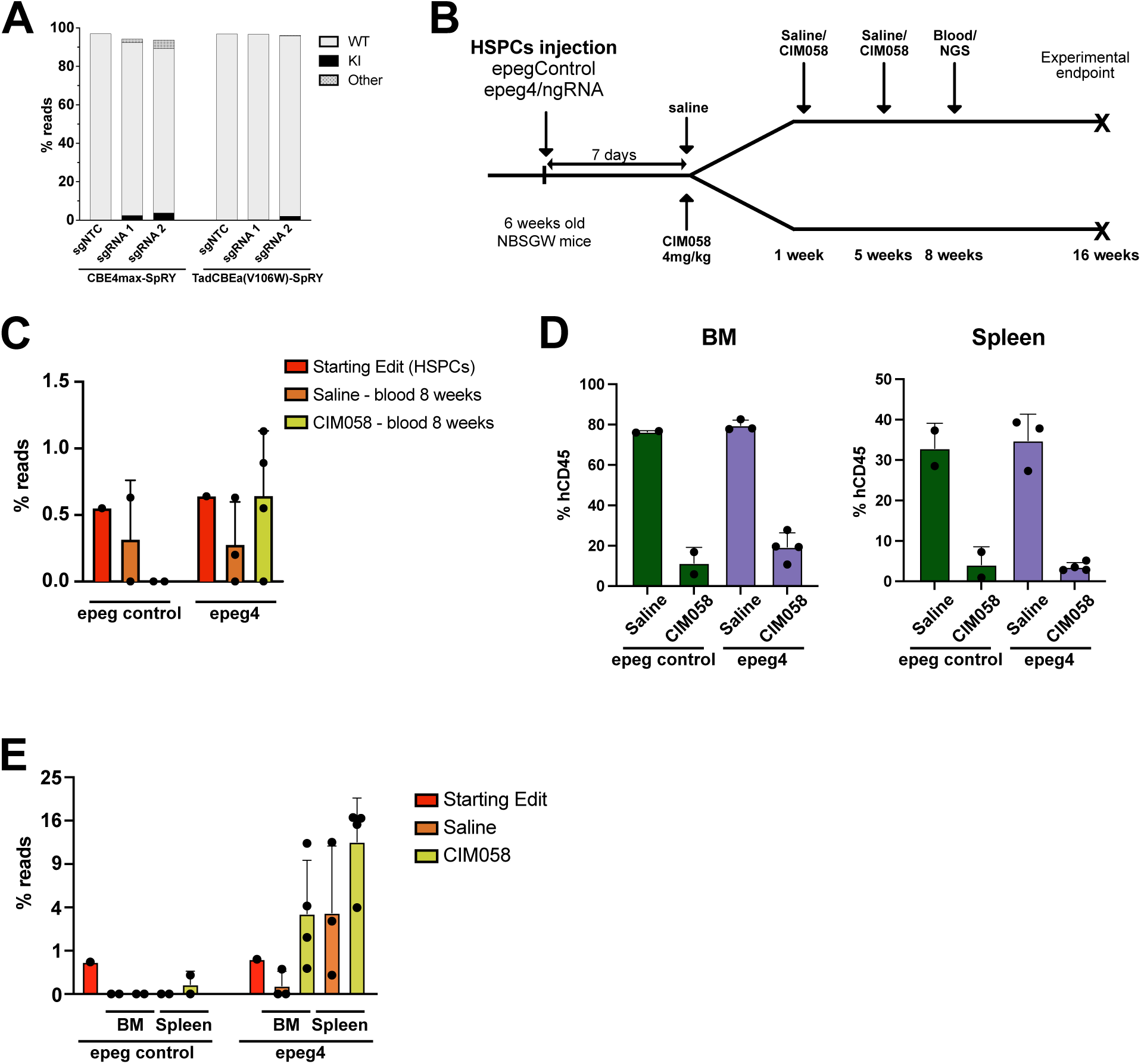
related to Figure 4: CD117^E73Kepeg4-PE^ HSPCs retained function and can be enriched in vivo (A) Quantification of the alleles by NGS. KI = E73K; Other = bystander edits (D72N; E73Stop; D72N + E73K; E73K + E76K; L71L + D72N + E73K; D72N + E73K + E76K) (B) Schematic representation of the mouse in vivo experiment. NBSGW mice were engrafted with epegRNA control or E73K epegRNA4 edited HSPCs and euthanised after 16 weeks. Mice received 3 injections of saline or CIM058 4mg/kg. (C) NGS analysis of the edited reads (KI for control: silent and PAM mutations; KI for E73K: E73K codon, silent and PAM mutations) in the blood of the mice 8 weeks after humanization. (D) FACS quantification of human CD45^+^ cells in the BM and the spleen of the mice at the end of the experiment. (E) NGS analysis of the edited reads (KI for control: silent and PAM mutations; KI for E73K: E73K codon, silent and PAM mutations) in the BM and the spleen of the mice at the end of the experiment.

**Figure S3:**
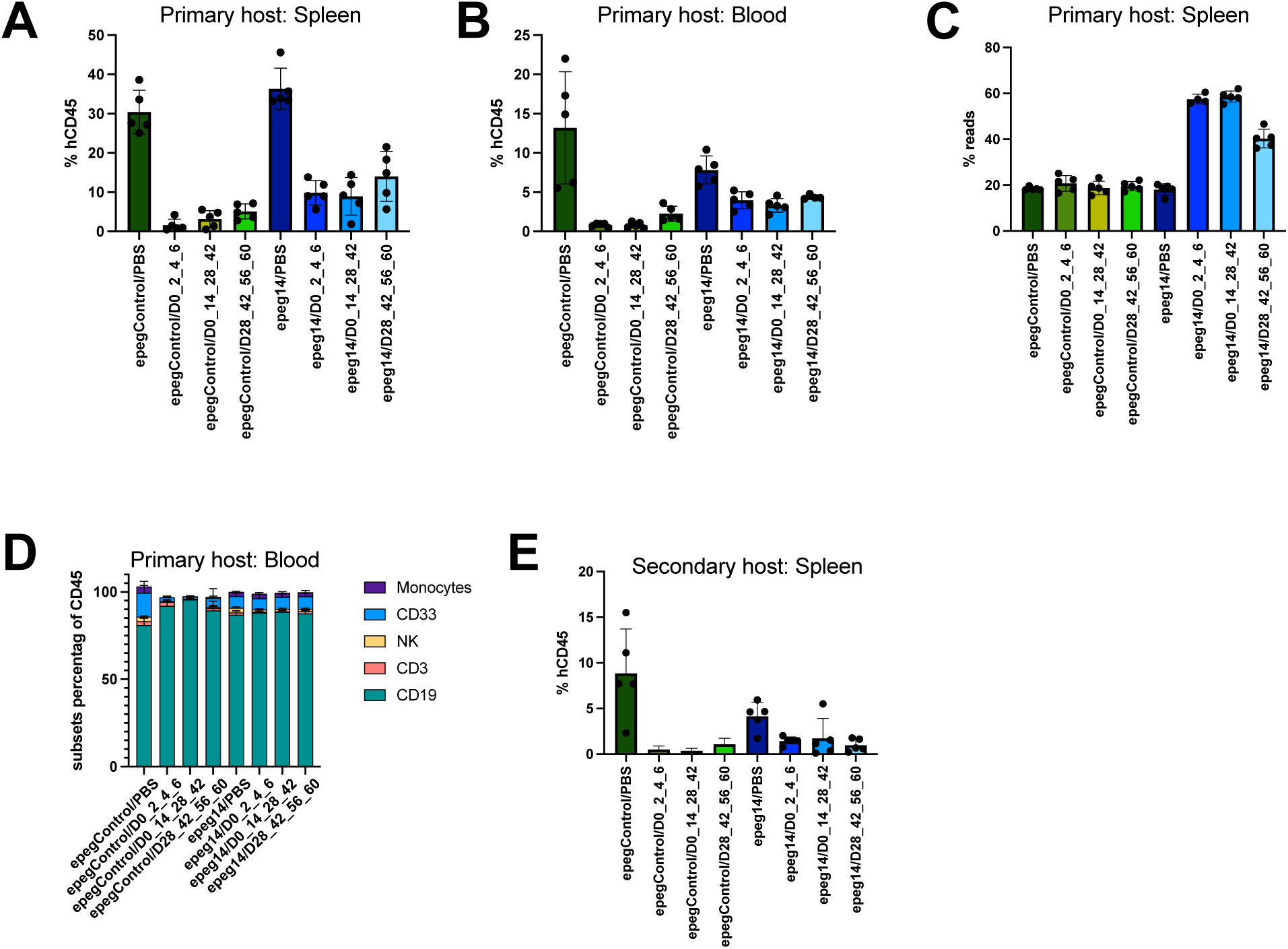
related to Figure 5: CD117^E73Kepeg14-PE^ HSPCs retained function and can be enriched in vivo (A-B) FACS quantification of human CD45^+^ cells in the spleen (A) and blood (B) of the mice at the end of the experiment. (D) NGS analysis of the edited reads (KI for control: silent and PAM mutations; KI for E73K: E73K codon, silent and PAM mutations) in the spleen of the mice at the end of the experiment. (E) Flow cytometry analysis of the multi-lineage differentiation in the blood of the mice at the end of the experiment. (F) FACS quantification of human CD45^+^ cells in the spleen of the mice at the end of secondary transplant experiment.

## REFERENCES

1. Copelan, E.A. (2006). Hematopoietic stem-cell transplantation. N Engl J Med 354, 1813–1826. 10.1056/NEJMra052638.

2. Gratwohl, A., Pasquini, M.C., Aljurf, M., Atsuta, Y., Baldomero, H., Foeken, L., Gratwohl, M., Bouzas, L.F., Confer, D., Frauendorfer, K., et al. (2015). One million haemopoietic stem-cell transplants: a retrospective observational study. Lancet Haematol 2, e91–100. 10.1016/S2352-3026(15)00028-9.

3. Kohn, D.B., Booth, C., Shaw, K.L., Xu-Bayford, J., Garabedian, E., Trevisan, V., Carbonaro-Sarracino, D.A., Soni, K., Terrazas, D., Snell, K., et al. (2021). Autologous Ex Vivo Lentiviral Gene Therapy for Adenosine Deaminase Deficiency. N Engl J Med 384, 2002–2013. 10.1056/NEJMoa2027675.

4. Kohn, D.B., and Candotti, F. (2009). Gene therapy fulfilling its promise. N Engl J Med 360, 518-521. 10.1056/NEJMe0809614.

5. Morgan, R.A., Gray, D., Lomova, A., and Kohn, D.B. (2017). Hematopoietic Stem Cell Gene Therapy: Progress and Lessons Learned. Cell Stem Cell 21, 574–590. 10.1016/j.stem.2017.10.010.

6. Frangoul, H., Altshuler, D., Cappellini, M.D., Chen, Y.S., Domm, J., Eustace, B.K., Foell, J., de la Fuente, J., Grupp, S., Handgretinger, R., et al. (2021). CRISPR-Cas9 Gene Editing for Sickle Cell Disease and beta-Thalassemia. N Engl J Med 384, 252–260. 10.1056/NEJMoa2031054.

7. Kanter, J., Walters, M.C., Krishnamurti, L., Mapara, M.Y., Kwiatkowski, J.L., Rifkin-Zenenberg, S., Aygun, B., Kasow, K.A., Pierciey, F.J., Jr., Bonner, M., et al. (2022). Biologic and Clinical Efficacy of LentiGlobin for Sickle Cell Disease. N Engl J Med 386, 617–628. 10.1056/NEJMoa2117175.

8. Booth, C., Masiuk, K., Vazouras, K., Fernandes, A., Xu-Bayford, J., Campo Fernandez, B., Roy, S., Curio-Penny, B., Arnold, J., Terrazas, D., et al. (2025). Long-Term Safety and Efficacy of Gene Therapy for Adenosine Deaminase Deficiency. N Engl J Med 393, 1486–1497. 10.1056/NEJMoa2502754.

9. John, T., and Czechowicz, A. (2025). Clinical hematopoietic stem cell-based gene therapy. Mol Ther 33, 2663–2678. 10.1016/j.ymthe.2025.04.029.

10. Farmakis, D., Porter, J., Taher, A., Domenica Cappellini, M., Angastiniotis, M., and Eleftheriou, A. (2022). 2021 Thalassaemia International Federation Guidelines for the Management of Transfusion-dependent Thalassemia. Hemasphere 6, e732. 10.1097/HS9.0000000000000732.

11. Higgs, D.R., Engel, J.D., and Stamatoyannopoulos, G. (2012). Thalassaemia. Lancet 379, 373–383. 10.1016/S0140-6736(11)60283-3.

12. Tuysuz, G., and Tayfun, F. (2017). Health-related Quality of Life and its Predictors Among Transfusion-dependent Thalassemia Patients. J Pediatr Hematol Oncol 39, 332–336. 10.1097/MPH.0000000000000790.

13. Jinek, M., Chylinski, K., Fonfara, I., Hauer, M., Doudna, J.A., and Charpentier, E. (2012). A programmable dual-RNA-guided DNA endonuclease in adaptive bacterial immunity. Science 337, 816–821. 10.1126/science.1225829.

14. Komor, A.C., Kim, Y.B., Packer, M.S., Zuris, J.A., and Liu, D.R. (2016). Programmable editing of a target base in genomic DNA without double-stranded DNA cleavage. Nature 533, 420–424. 10.1038/nature17946.

15. Gaudelli, N.M., Komor, A.C., Rees, H.A., Packer, M.S., Badran, A.H., Bryson, D.I., and Liu, D.R. (2017). Programmable base editing of A*T to G*C in genomic DNA without DNA cleavage. Nature 551, 464–471. 10.1038/nature24644.

16. Anzalone, A.V., Randolph, P.B., Davis, J.R., Sousa, A.A., Koblan, L.W., Levy, J.M., Chen, P.J., Wilson, C., Newby, G.A., Raguram, A., and Liu, D.R. (2019). Search-and-replace genome editing without double-strand breaks or donor DNA. Nature 576, 149–157. 10.1038/s41586-019-1711-4.

17. Cong, L., Ran, F.A., Cox, D., Lin, S., Barretto, R., Habib, N., Hsu, P.D., Wu, X., Jiang, W., Marraffini, L.A., and Zhang, F. (2013). Multiplex genome engineering using CRISPR/Cas systems. Science 339, 819–823. 10.1126/science.1231143.

18. Mali, P., Yang, L., Esvelt, K.M., Aach, J., Guell, M., DiCarlo, J.E., Norville, J.E., and Church, G.M. (2013). RNA-guided human genome engineering via Cas9. Science 339, 823–826. 10.1126/science.1232033.

19. Frangoul, H., Locatelli, F., Sharma, A., Bhatia, M., Mapara, M., Molinari, L., Wall, D., Liem, R.I., Telfer, P., Shah, A.J., et al. (2024). Exagamglogene Autotemcel for Severe Sickle Cell Disease. N Engl J Med 390, 1649–1662. 10.1056/NEJMoa2309676.

20. Gupta, A.O., Sharma, A., Frangoul, H., Kanter, J., Mapara, M.Y., Dalal, J., Alavi, A., Jaroscak, J.J., Ayala, E., DiPersio, J.F., et al. (2026). Base Editing of HBG1 and HBG2 Promoters for Sickle Cell Disease. N Engl J Med 394, 1824–1835. 10.1056/NEJMoa2504835.

21. Gori, J.L., Haddad, E., Frangoul, H., Kohn, D.B., Morris, E.C., Martin, B.N., Deary, B.A., Nickerson, M., Scholz, R.L., Fernandez, I., et al. (2026). Prime Editing for p47(phox)-Deficient Chronic Granulomatous Disease. N Engl J Med 394, 1195–1203. 10.1056/NEJMoa2509807.

22. Finck, A.V., Blanchard, T., Roselle, C.P., Golinelli, G., and June, C.H. (2022). Engineered cellular immunotherapies in cancer and beyond. Nat Med 28, 678–689. 10.1038/s41591-022-01765-8.

23. McCune, J.M., and Kiem, H.P. (2024). Extending Gene Medicines to All in Need. N Engl J Med 390, 1721–1722. 10.1056/NEJMe2403104.

24. Rouce, R.H., and Porteus, M.H. (2024). Cell and gene therapy accessibility. Science 385, 475. 10.1126/science.ads0252.

25. Gyurkocza, B., and Sandmaier, B.M. (2014). Conditioning regimens for hematopoietic cell transplantation: one size does not fit all. Blood 124, 344–353. 10.1182/blood-2014-02-514778.

26. Uchida, N., Stasula, U., Demirci, S., Germino-Watnick, P., Hinds, M., Le, A., Chu, R., Berg, A., Liu, X., Su, L., et al. (2023). Fertility-preserving myeloablative conditioning using single-dose CD117 antibody-drug conjugate in a rhesus gene therapy model. Nat Commun 14, 6291. 10.1038/s41467-023-41153-5.

27. Long-Boyle, J.R., Kohn, D.B., Shah, A.J., Spencer, S.M., Sevilla, J., Booth, C., Lopez Lorenzo, J.L., Nicoletti, E., Shah, A., Reatz, M., et al. (2024). Busulfan and subsequent malignancy: An evidence-based risk assessment. Pediatr Blood Cancer 71, e30738. 10.1002/pbc.30738.

28. Faraci, M., Diesch, T., Labopin, M., Dalissier, A., Lankester, A., Gennery, A., Sundin, M., Uckan-Cetinkaya, D., Bierings, M., Peters, A.M.J., et al. (2019). Gonadal Function after Busulfan Compared with Treosulfan in Children and Adolescents Undergoing Allogeneic Hematopoietic Stem Cell Transplant. Biol Blood Marrow Transplant 25, 1786–1791. 10.1016/j.bbmt.2019.05.005.

29. Frangoul, H., de la Fuente, J., Chopra, Y., Meisel, R., Amrolia, P.J., Algeri, M., Sharma, A., Cappellini, M.D., Corbacioglu, S., Kattamis, A., et al. (2026). Exa-cel in Children with Transfusion-Dependent beta-Thalassemia or Sickle Cell Disease. N Engl J Med. 10.1056/NEJMoa2603387.

30. Czechowicz, A., Kraft, D., Weissman, I.L., and Bhattacharya, D. (2007). Efficient transplantation via antibody-based clearance of hematopoietic stem cell niches. Science 318, 1296–1299. 10.1126/science.1149726.

31. Pang, W.W., Czechowicz, A., Logan, A.C., Bhardwaj, R., Poyser, J., Park, C.Y., Weissman, I.L., and Shizuru, J.A. (2019). Anti-CD117 antibody depletes normal and myelodysplastic syndrome human hematopoietic stem cells in xenografted mice. Blood 133, 2069–2078. 10.1182/blood-2018-06-858159.

32. Kwon, H.S., Logan, A.C., Chhabra, A., Pang, W.W., Czechowicz, A., Tate, K., Le, A., Poyser, J., Hollis, R., Kelly, B.V., et al. (2019). Anti-human CD117 antibody-mediated bone marrow niche clearance in nonhuman primates and humanized NSG mice. Blood 133, 2104–2108. 10.1182/blood-2018-06-853879.

33. Agarwal, R., Dvorak, C.C., Prockop, S., Kwon, H.S., Long-Boyle, J.R., Le, A., Brown, J.W., Merkel, E., Truong, K., Velasco, B., et al. (2021). JSP191 As a Single-Agent Conditioning Regimen Results in Successful Engraftment, Donor Myeloid Chimerism, and Production of Donor Derived Naïve Lymphocytes in Patients with Severe Combined Immunodeficiency (SCID). Blood 138 (*1*). 10.1182/blood-2021-153074.

34. Jung, D., Long-Boyle, J.R., Pang, W.W., and Gobburu, J.V.S. (2024). Pharmacokinetics of Briquilimab as a Conditioning Agent for Hematopoietic Stem Cell Transplantation in Patients With Severe Combined Immunodeficiency, Myelodysplastic Syndrome, or Acute Myeloid Leukemia. Transplant Cell Ther 30, 923 e921–923 e929. 10.1016/j.jtct.2024.07.001.

35. Agarwal, R.K., Dvorak, C.C., Kwon, H.-S., Long-Boyle, J.R., Prohaska, S.S., Brown, J.W., Le, A., Guttman-Klein, A., Weissman, I.L., Cowan, M.J., et al. (2019). Non-genotoxic anti-CD117 antibody conditioning results in successful hematopoietic stem cell engraftment in patients with severe combined immunodeficiency. Blood 134. 10.1182/blood-2019-126239.

36. Agarwal, R., Bertaina, A., Soco, C., Long-Boyle, J.R., Saini, G., Kunte, N., Hiroshima, L., Chan, Y.Y., Willner, H., Krampf, M.R., et al. (2025). Irradiation- and busulfan-free stem cell transplantation in Fanconi anemia using an anti-CD117 antibody: a phase 1b trial. Nat Med 31, 3183–3190. 10.1038/s41591-025-03817-1.

37. Garaude, S., Marone, R., Lepore, R., Devaux, A., Beerlage, A., Seyres, D., Dell’ Aglio, A., Juskevicius, D., Zuin, J., Burgold, T., et al. (2024). Selective haematological cancer eradication with preserved haematopoiesis. Nature 630, 728–735. 10.1038/s41586-024-07456-3.

38. Marone, R., Landmann, E., Devaux, A., Lepore, R., Seyres, D., Zuin, J., Burgold, T., Engdahl, C., Capoferri, G., Dell’Aglio, A., et al. (2023). Epitope-engineered human hematopoietic stem cells are shielded from CD123-targeted immunotherapy. J Exp Med 220. 10.1084/jem.20231235.

39. Casirati, G., Cosentino, A., Mucci, A., Salah Mahmoud, M., Ugarte Zabala, I., Zeng, J., Ficarro, S.B., Klatt, D., Brendel, C., Rambaldi, A., et al. (2023). Epitope editing enables targeted immunotherapy of acute myeloid leukaemia. Nature 621, 404–414. 10.1038/s41586-023-06496-5.

40. Wellhausen, N., O’Connell, R.P., Lesch, S., Engel, N.W., Rennels, A.K., Gonzales, D., Herbst, F., Young, R.M., Garcia, K.C., Weiner, D., et al. (2023). Epitope base editing CD45 in hematopoietic cells enables universal blood cancer immune therapy. Sci Transl Med 15, eadi1145. 10.1126/scitranslmed.adi1145.

41. Davidson, E., and Doranz, B.J. (2014). A high-throughput shotgun mutagenesis approach to mapping B-cell antibody epitopes. Immunology 143, 13–20. 10.1111/imm.12323.

42. Reshetnyak, A.V., Nelson, B., Shi, X., Boggon, T.J., Pavlenco, A., Mandel-Bausch, E.M., Tome, F., Suzuki, Y., Sidhu, S.S., Lax, I., and Schlessinger, J. (2013). Structural basis for KIT receptor tyrosine kinase inhibition by antibodies targeting the D4 membrane-proximal region. Proc Natl Acad Sci U S A 110, 17832–17837. 10.1073/pnas.1317118110.

43. L’Italien, L., Orozco, O., Abrams, T., Cantagallo, L., Connor, A., Desai, J., Ebersbach, H., Gelderblom, H., Hoffmaster, K., Lees, E., et al. (2018). Mechanistic Insights of an Immunological Adverse Event Induced by an Anti-KIT Antibody Drug Conjugate and Mitigation Strategies. Clin Cancer Res 24, 3465–3474. 10.1158/1078-0432.CCR-17-3786.

44. Turchiano, G., Andrieux, G., Klermund, J., Blattner, G., Pennucci, V., El Gaz, M., Monaco, G., Poddar, S., Mussolino, C., Cornu, T.I., et al. (2021). Quantitative evaluation of chromosomal rearrangements in gene-edited human stem cells by CAST-Seq. Cell Stem Cell 28, 1136–1147 e1135. 10.1016/j.stem.2021.02.002.

45. Koblan, L.W., Doman, J.L., Wilson, C., Levy, J.M., Tay, T., Newby, G.A., Maianti, J.P., Raguram, A., and Liu, D.R. (2018). Improving cytidine and adenine base editors by expression optimization and ancestral reconstruction. Nat Biotechnol 36, 843–846. 10.1038/nbt.4172.

46. Neugebauer, M.E., Hsu, A., Arbab, M., Krasnow, N.A., McElroy, A.N., Pandey, S., Doman, J.L., Huang, T.P., Raguram, A., Banskota, S., et al. (2023). Evolution of an adenine base editor into a small, efficient cytosine base editor with low off-target activity. Nat Biotechnol 41, 673–685. 10.1038/s41587-022-01533-6.

47. Chen, P.J., Hussmann, J.A., Yan, J., Knipping, F., Ravisankar, P., Chen, P.F., Chen, C., Nelson, J.W., Newby, G.A., Sahin, M., et al. (2021). Enhanced prime editing systems by manipulating cellular determinants of editing outcomes. Cell 184, 5635–5652 e5629. 10.1016/j.cell.2021.09.018.

48. Yannaki, E., Papayannopoulou, T., Jonlin, E., Zervou, F., Karponi, G., Xagorari, A., Becker, P., Psatha, N., Batsis, I., Kaloyannidis, P., et al. (2012). Hematopoietic stem cell mobilization for gene therapy of adult patients with severe beta-thalassemia: results of clinical trials using G-CSF or plerixafor in splenectomized and nonsplenectomized subjects. Mol Ther 20, 230–238. 10.1038/mt.2011.195.

49. Czechowicz, A., Palchaudhuri, R., Scheck, A., Hu, Y., Hoggatt, J., Saez, B., Pang, W.W., Mansour, M.K., Tate, T.A., Chan, Y.Y., et al. (2019). Selective hematopoietic stem cell ablation using CD117-antibody-drug-conjugates enables safe and effective transplantation with immunity preservation. Nat Commun 10, 617. 10.1038/s41467-018-08201-x.

50. Alvarado, D., Maurer, M., Gedrich, R., Seibel, S.B., Murphy, M.B., Crew, L., Goldstein, J., Crocker, A., Vitale, L.A., Morani, P.A., et al. (2022). Anti-KIT monoclonal antibody CDX-0159 induces profound and durable mast cell suppression in a healthy volunteer study. Allergy 77, 2393–2403. 10.1111/all.15262.

51. Metz, M., Mitha, E., Leflein, J., Talreja, N., Gotua, M., Krasowska, D., Peter, J., Anderson, J., Young, D., Heath-Chiozzi, M., et al. (2026). Randomized dose-finding study of anti-KIT barzolvolimab in patients with chronic spontaneous urticaria. J Allergy Clin Immunol. 10.1016/j.jaci.2026.02.018.

52. Agarwal, R., Weinberg, K.I., Kwon, H.S., Le, A., Long-Boyle, J.R., Kohn, D.B., Bradford, K., De Oliveira, S., Bertaina, A., Czechowicz, A., et al. (2020). First Report of Non-Genotoxic Conditioning with JSP191 (anti-CD117) and Hematopoietic Stem Cell Transplantation in a Newly Diagnosed Patient with Severe Combined Immune Deficiency. Blood 136 (*1*). 10.1182/blood-2020-137762.

53. Maurer, M., Metz, M., Anderson, J., Talreja, N., Young, D., Crowley, E., Heath-Chiozzi, M., Ma, R., Paradise, E., Hawthorne, T., et al. (2025). Anti-KIT Barzolvolimab for Chronic Spontaneous Urticaria. Allergy 80, 2178–2186. 10.1111/all.16598.

54. Paul, S., Konig, M.F., Pardoll, D.M., Bettegowda, C., Papadopoulos, N., Wright, K.M., Gabelli, S.B., Ho, M., van Elsas, A., and Zhou, S. (2024). Cancer therapy with antibodies. Nat Rev Cancer 24, 399–426. 10.1038/s41568-024-00690-x.

55. DiPersio, J.F., Koehne, G., Shah, N.N., Bernard, L., Suh, H.C., Koura, D., Tamari, R., Mushtaq, M.U., Maakaron, J., Rimando, J., et al. (2026). CRISPR-Cas9 CD33-deleted allogeneic hematopoietic cell transplantation with gemtuzumab ozogamicin maintenance in AML: a phase 1/2 trial. Nat Med 32, 1763–1772. 10.1038/s41591-026-04362-1.

56. Demirci, S., Mondal, N., Austin, W., Butt, H., London, E., Sathish, S., Zhang, K., Wong, J., Kromer, H., Harmon, A., et al. (2024). CD117 Antibody Conditioning and Multiplex Base Editing Enable Rapid and Robust Fetal Hemoglobin Reactivation in a Rhesus Autologous Transplantation Model. Blood 144. 10.1182/blood-2024-204403.

57. Sakai, H.A., Pierce, S.E., Jiang, A.Y., Cristian, A., An, M., Kim, C.R., Ahmed, N., Hemez, C.F., Tao, Y.A., Zhang, E., et al. (2026). Directed evolution of small RNA-stabilizing motifs that improve prime-editing efficiency. Nat Biotechnol. 10.1038/s41587-026-03123-2.

58. Heath, J.M., Tedeschi, J.G., Arvindam, U.S., Padhye, S., Laoharawee, K., Ng, A.C., Waterman, D.p., Roy, M.S., Alexander, S.C., Trusiak, S., et al. (2024). Prime Editing Enables Precise and Efficient Single Amino Acid Substitutions to Shield CD34+ Hematopoietic Stem Cells from Anti-CD117 Antibody-Based Conditioning. Blood 144 (*1*), 514. 10.1182/blood-2024-203602.

59. https://primemedicine.com/wp-content/uploads/2025/10/ASH-CD34-HSC-Shielding-Presentation.pdf.

60. Corbacioglu, S., Troeger, A., Kleinschmidt, K., Hanafee-Alali, T., Brosig, A.-M., Jakob, M., Kramer, S., Offner, R., Wolff, D., and Foell, J. (2023). Haploidentical Aß T Cell Depleted HSCT Represents an Alternative Treatment Option in Pediatric and Adult Patients with Sickle Cell Disease (SCD). Blood 142 (*1*). 10.1182/blood-2023-189434.

61. Li, Z., and Murphy, P.M. (2022). CD45: a niche marker for allotransplantation. Blood 139 (*1*). 10.1182/blood.2021015024.

62. Cao, A., and Galanello, R. (2010). Beta-thalassemia. Genet Med 12, 61–76. 10.1097/GIM.0b013e3181cd68ed.

63. https://www.fda.gov/news-events/press-announcements/fda-issues-draft-guidance-genome-editing-safety-standards-advance-gene-therapy-development.

64. https://www.fda.gov/regulatory-information/search-fda-guidance-documents/safety-assessment-genome-editing-human-gene-therapy-products-using-next-generation-sequencing.

65. Kluesner, M.G., Tasakis, R.N., Lerner, T., Arnold, A., Wust, S., Binder, M., Webber, B.R., Moriarity, B.S., and Pecori, R. (2021). MultiEditR: The first tool for the detection and quantification of RNA editing from Sanger sequencing demonstrates comparable fidelity to RNA-seq. Mol Ther Nucleic Acids 25, 515–523. 10.1016/j.omtn.2021.07.008.

66. https://github.com/ivanek/MultiEditR.

67. Clement, K., Rees, H., Canver, M.C., Gehrke, J.M., Farouni, R., Hsu, J.Y., Cole, M.A., Liu, D.R., Joung, J.K., Bauer, D.E., and Pinello, L. (2019). CRISPResso2 provides accurate and rapid genome editing sequence analysis. Nat Biotechnol 37, 224–226. 10.1038/s41587-019-0032-3.

68. Rhiel, M., Geiger, K., Andrieux, G., Rositzka, J., Boerries, M., Cathomen, T., and Cornu, T.I. (2023). T-CAST: An optimized CAST-Seq pipeline for TALEN confirms superior safety and efficacy of obligate-heterodimeric scaffolds. Front Genome Ed 5, 1130736. 10.3389/fgeed.2023.1130736.

69. Giarratana, M.C., Rouard, H., Dumont, A., Kiger, L., Safeukui, I., Le Pennec, P.Y., Francois, S., Trugnan, G., Peyrard, T., Marie, T., et al. (2011). Proof of principle for transfusion of in vitro-generated red blood cells. Blood 118, 5071–5079. 10.1182/blood-2011-06-362038.

70. Bankhead, P., Loughrey, M.B., Fernandez, J.A., Dombrowski, Y., McArt, D.G., Dunne, P.D., McQuaid, S., Gray, R.T., Murray, L.J., Coleman, H.G., et al. (2017). QuPath: Open source software for digital pathology image analysis. Sci Rep 7, 16878. 10.1038/s41598-017-17204-5.

